# New Human IPSC Models of Late-onset Alzheimer’s Disease Polygenic Risk Identify Multiple Impairments of Microglial Function

**DOI:** 10.64898/2026.04.29.721607

**Authors:** Hazel Hall-Roberts, Emily Maguire, Bethany Shaw, Rachel O’Donoghue, Jincy Winston, Samuel Keat, Lorenzo Capitani, Rebecca Mahoney, Bethany Geary, Karolina Faber, Nina Stöberl, Natalie Connor-Robson, Charlotte Bridge, Atahualpa Castillo Morales, Mateus Bernado-Harrington, Nicholas D. Allen, Valentina Escott-Price, Caleb Webber, Sally A. Cowley, Philip R. Taylor, Peter Holmans, Rebecca Sims, Julie Williams

**Affiliations:** UK Dementia Research Institute at Cardiff University, Maindy Road, Cardiff CF24 4HQ, UK; MRC Protein Phosphorylation and Ubiquitylation Unit, School of Life Sciences, University of Dundee, Dundee DD1 4HN, UK; James and Lillian Martin Centre for Stem Cell Research, Sir William Dunn School of Pathology, University of Oxford, South Parks Road, Oxford OX1 3RE, UK; School of Biosciences, Cardiff University, Museum Avenue, Cardiff CF10 3AX, UK; Division of Psychological Medicine and Clinical Neurosciences, Cardiff University, Maindy Road, Cardiff CF24 4HQ, UK; Moondance Dementia Laboratory, Maindy Road, Cardiff, CF24 4HQ, UK; Systems Immunity Research Institute, Cardiff University, Heath Park, Cardiff CF14 4XN, UK

**Author notes:** Joint first authors (Hazel Hall-Roberts, Emily Maguire). Joint corresponding authors (Hazel Hall-Roberts, Julie Williams). Centre for Medicines Discovery, Nuffield Department of Medicine, NDM Research Building, Old Road Campus, Oxford, OX3 7FZ.

## Abstract

Genetic discoveries implicate microglia in late-onset Alzheimer’s disease (AD). We modelled AD in a powerful study of 51 human induced pluripotent stem cell (iPSC) microglia derived from high-polygenic risk AD or low-risk cognitively well individuals, sampled from a large cohort. We explored mitochondrial function, cytokine secretion, endocytosis, phagocytosis, lipid accumulation, calcium store release, and chemotaxis under basal conditions and immune challenge. High polygenic risk was independently associated with significant functional deficits in iPSC microglia under immune challenge, in mitochondrial ATP production (p=0.005, −13%), and inflammatory cytokine release (IL-6: p=0.018, −42%; TNF p=0.026, −38.5%). Furthermore, a selective deficit in amyloid-β uptake was identified (p=0.00477, −5.9%). Deficits in inflammatory cytokine release were driven by *APOE* ε4. These findings reflect primary changes in AD pathogenesis predating plaque formation and validate a human *in vitro* platform for late-onset AD to further understand disease mechanisms and screen drug or genetic therapies.

## Introduction

Alzheimer’s disease (AD) is the leading cause of dementia worldwide, affecting tens of millions of people and placing a substantial burden on individuals, families, and healthcare systems. The majority of cases (∼95%) occur after the age of 65 (late-onset AD, LOAD) (Reitz et al., 2020). However, modelling this common form of AD is challenging. LOAD arises from a complex interplay of genetic and environmental factors, with heritability estimated at 60–80% (Gatz et al., 2006). Genome-wide association studies (GWAS) have currently identified over 90 risk loci contributing to disease susceptibility (ADGC et al., 2025). These implicate several potential mechanistic pathways, including different aspects of the brain’s immune system, endocytosis and lipid processing, that may converge on the causal pathway for LOAD (Sims et al., 2020). We, and others, have captured this combined risk in a polygenic risk score, which has a positive predictive value for AD of 80% in pathologically confirmed cases, rising to over 90% accuracy at the polygenic-risk extremes (Escott-Price et al., 2015; Leonenko et al., 2021). Our strategy was to create a powerful sample of human induced pluripotent stem cells (iPSC) reflecting polygenic extremes, enabling us to model common LOAD *in vitro*, to explore disease mechanisms, and provide a platform for future therapeutic testing.

Genetic evidence strongly implicates microglia in AD pathology, with many of the LOAD-associated GWAS variants found within loci containing genes highly or exclusively expressed in microglia, or in non-coding regulatory regions active in these cells (Efthymiou & Goate, 2017; Tansey et al., 2018). Studies of single variants have shed light on how individual risk alleles affect microglial function, but they do not capture the polygenic nature of LOAD (De Deyn & Sleegers, 2025; Hall-Roberts et al., 2020; Maguire et al., 2021). Each person may carry many variants that together influence risk, most with only very small individual effects. iPSC technology enables the generation of microglia-like cells that retain donor genotypes, providing a powerful platform to study how cumulative genetic risk shapes microglial biology in a way that reflects real-world populations. In the healthy brain, microglia perform essential protective functions, including cytokine and chemokine release, phagocytosis, and endocytosis. In response to an immune challenge, microglia upregulate these immune functions, which all require a substantial amount of energy in the form of adenosine triphosphate (ATP) (Jung et al., 2025). Microglia can dynamically reprogram their metabolism to generate the ATP required to upregulate specific immune functions, thus enabling them to respond effectively to changing brain environments (Jung et al., 2025).

We aimed to uncover functional pathways in microglia altered by polygenic risk for AD, including those triggered by immune challenge where appropriate. Our study utilised a cohort of over 8,000 well characterised AD cases and controls, from which we derived iPSC lines from 51 individuals with extremes of polygenic AD risk, either high-risk LOAD (HRLOAD), or low-risk aged cognitively well controls (LRWELL) (Maguire et al., 2025). We transformed these iPSC into microglia-like cells and passed them through a comprehensive battery of assays that assessed disease relevant, functional pathways including measures of mitochondrial function, inflammatory cytokine secretion, lipid accumulation, calcium store release, endocytic uptake, phagocytosis, and chemotaxis (Maguire et al., 2021, 2022; Maninger et al., 2024; Mazaheri et al., 2017; Muth et al., 2019; Piers et al., 2019; Victor et al., 2022). In addition, we modelled immune challenge, stimulating cells with lipopolysaccharide (LPS), which has been shown to induce a transcriptional phenotype resembling that observed in AD mouse models, in iPSC-derived microglia (Monzón-Sandoval et al., 2022).

By comparing functional pathways of HRLOAD with LRWELL, we observed altered microglia functions, including significant deficits in mitochondrial metabolic reprogramming, inflammatory cytokine secretion under immune challenge, and a selective deficit in basal amyloid-β uptake. Our findings advance current understanding of how high LOAD genetic risk shapes microglial biology. This study provides a reproducible framework that can be applied to other iPSC-derived cell types to further dissect AD mechanisms, and validates a novel *in vitro* platform of common forms of LOAD for testing emerging therapeutics.

## Results

We differentiated 34 HRLOAD and 17 LRWELL iPSC lines (Supplementary Table 1) into microglia across six batches, performing functional assays and proteomics in parallel. High efficiency of differentiation was achieved (94% IBA1+, Extended Data Figure 2). Immune challenge was modelled, where appropriate, in our analyses comparing HRLOAD with LRWELL. Statistical analysis included correction for multiple testing and investigated significant associations in the effects of polygenic risk score on outcomes with pairwise comparisons (all corrected and uncorrected *p*-values are listed in Supplementary Table 2). We also tested significant differences based on the presence or absence of *APOE* ε4 alleles, where appropriate, and assessed relationships between significant findings using analysis of correlations between them.

### High polygenic risk iPSC microglia exhibit impaired metabolic reprogramming

Microglia respond quickly to changes in their environment, for example by producing inflammatory mediators in response to immune challenge. To fuel rapid physiological changes, microglia exhibit metabolic flexibility. They quickly upregulate glycolytic ATP production via metabolic reprogramming, temporarily favouring this more rapid form of energy production over the slower (but more efficient) oxidative phosphorylation (Miao et al., 2023). Immune challenge of microglia was induced using LPS, with ATP production examined using a Seahorse XF ATP Rate Assay (Fig 1A-D, Extended Data Figure 1). We observed a significant attenuation effect of HRLOAD on total ATP production only in the presence of LPS. Statistical analysis included correction for multiple testing and investigated significant differences with pairwise comparisons stimulation, indicated by the HRLOAD:LPS interaction (Fig 1A, uncorrected p= 5.4 × 10^-4^, corrected p= 3.6 × 10^-3^, −13%). Underlying this was lower glycolytic ATP production (Fig 1B, uncorrected p=0.011, corrected p=0.033, −0.3%), and an absolute drop in mitochondrial ATP production (Fig 1C, uncorrected p=0.00096, corrected p=0.005, −13.4%) in HRLOAD challenged with LPS, versus challenged LRWELL. LRWELL microglia responded to LPS challenge by metabolic reprogramming, boosting glycolytic ATP production (Fig 1B) to elevate total ATP production rates (Fig 1A), with no change to mitochondrial ATP production (Fig 1C). HRLOAD status had no effect on ATP production without immune challenge. To illustrate the effect of HRLOAD on metabolic reprogramming, we also present the data as a ratio of LPS-challenged to unstimulated (Fig 1D), showing HRLOAD microglia had a weaker fold-change increase in glycolytic and total ATP production than LRWELL, and a reduction in mitochondrial ATP production. *APOE ε4* status had no significant effect on polygenic risk or LPS-induced changes to ATP production rate (Extended Data Figure 1).

**Figure 1.**
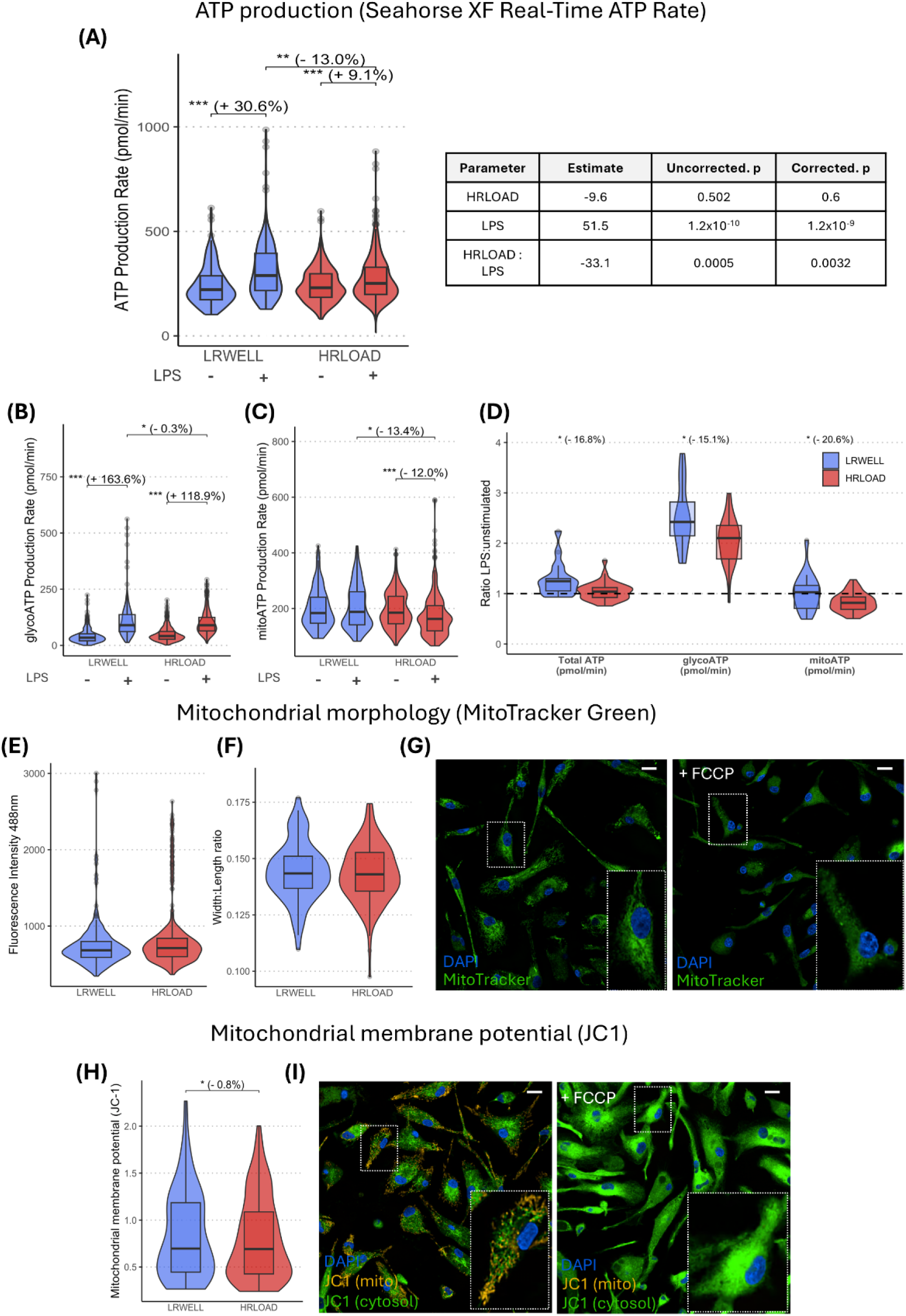
HRLOAD microglia exhibit mitochondrial metabolic dysfunction. (**A**) Total ATP production rate measured in the Seahorse XF Real-Time ATP Rate assay, with and without LPS stimulation (20ng/mL for 18 hours). Table shows results from the linear mixed effects model, including corrected and uncorrected p values. (**B**) ATP production rate from glycolysis (glycoATP) and (**C**) ATP production rate from mitochondrial respiration (mitoATP) measured in the Seahorse XF Real-Time ATP Rate assay, with and without LPS challenge (20ng/mL for 18 hours). (**D**) Fold change of the LPS-challenged ATP Rate assay data relative to unstimulated (raw data presented in (**A-C**)). For (**A-D**), data was first normalised to cell count, then a median obtained from 3-10 replicate wells for each experiments. (**E**) Total mitochondrial content (measured by mean fluorescence intensity of MitoTracker Green) and (**F**) mitochondrial network width:length ratio (measured using Mitotracker Green morphology) with representative images (± FCCP) shown in **(G)**. **(H)** Mitochondrial membrane potential analysed by JC1 red/green fluorescence intensity, with representative images (± FCCP) shown in **(I)**. For **(E, F and H)**, data plotted is mean per cell, then medians obtained from 4 replicate wells for each experiment. Across all parts of the figure, box plots show the median and interquartile range for N = 35 HRLOAD, 17 LRWELL from a minimum of 4 experiments. HRLOAD (red) was compared to LRWELL (blue), and LPS-challenged to unstimulated, using linear mixed effects modelling with p-values corrected by Benjamini-Hochberg procedure. * p < 0.05, ** p < 0.01, *** p < 0.001. Scale bars: 20 µm.

Mitochondria are central regulators of cellular metabolism and energy balance, processes that are particularly important for microglial activation and immune responses (Miao et al., 2023). Because changes in mitochondrial structure or function can strongly influence how microglia respond to stress (Jenkins et al., 2024), we compared mitochondrial features in LRWELL and HRLOAD iPSC-derived microglia without stimulation. We first assessed total mitochondrial mass (Fig 1E) and network morphology (Fig 1F), using MitoTracker Green (representative images and FCCP positive control shown in Fig 1G), which did not differ significantly between LRWELL and HRLOAD microglia. To assess the health of the mitochondrial network under basal conditions, we measured mitochondrial membrane potential (MMP) with JC-1 dye (Fig 1H). JC-1 forms red-fluorescent aggregates in healthy polarized mitochondria and remains green in the cytoplasm or depolarized mitochondria, therefore a lower red/green fluorescence ratio reflects reduced MMP (representative images and FCCP positive control shown in Fig 1I). HRLOAD microglia showed a slight, but significant, reduction in this ratio compared to LRWELL (Fig 1H, uncorrected p= 0.012, corrected p = 0.034, −0.8%), indicating decreased mitochondrial polarization.

Taken together, these findings show that HRLOAD microglia maintain normal ATP levels under resting conditions but have lower MMP and show a reduced ability to increase ATP production in response to metabolically demanding, immune challenged conditions.

### High polygenic risk iPSC microglia show impaired inflammatory cytokine responses

Inflammatory signalling is strongly implicated in AD progression (Chen et al., 2024), therefore we analysed a selection of secreted inflammation-associated proteins from our microglia under basal and immune challenged conditions using a LEGENDplex multiplex immunoassay, comprising sTREM2, IL-6, TNF, IL-10, CCL2, and IL-1β. Under immune challenge with LPS, HRLOAD significantly reduced the secretion of pro-inflammatory cytokines IL-6 (Fig 2A, uncorrected p= 4 × 10^-3^, corrected p= 0.016, −41.8%), and TNF (Fig 2C, uncorrected p= 7 × 10^-3^, corrected p= 0.026, −38.5%). Polygenic risk had no significant effect on secreted protein levels without stimulation (Supplementary Table 2), and did not significantly interact with LPS-challenged IL-10 (anti-inflammatory cytokine, Fig 2F), CCL2 (chemokine, Fig 2G), and IL-1β (pro-inflammatory cytokine, Fig 2H) secretion. The secreted fragment of the TREM2 receptor, sTREM2, trended towards a decrease with high polygenic risk at baseline (uncorrected p= 0.02, corrected p= 0.057; Fig 2E). When segregated by *APOE ε4* status, the reduction in IL-6 (Fig 2B) and TNF (Fig 2D) production following immune challenge was driven by HRLOAD lines carrying one or two *APOEε4* alleles.

**Figure 2.**
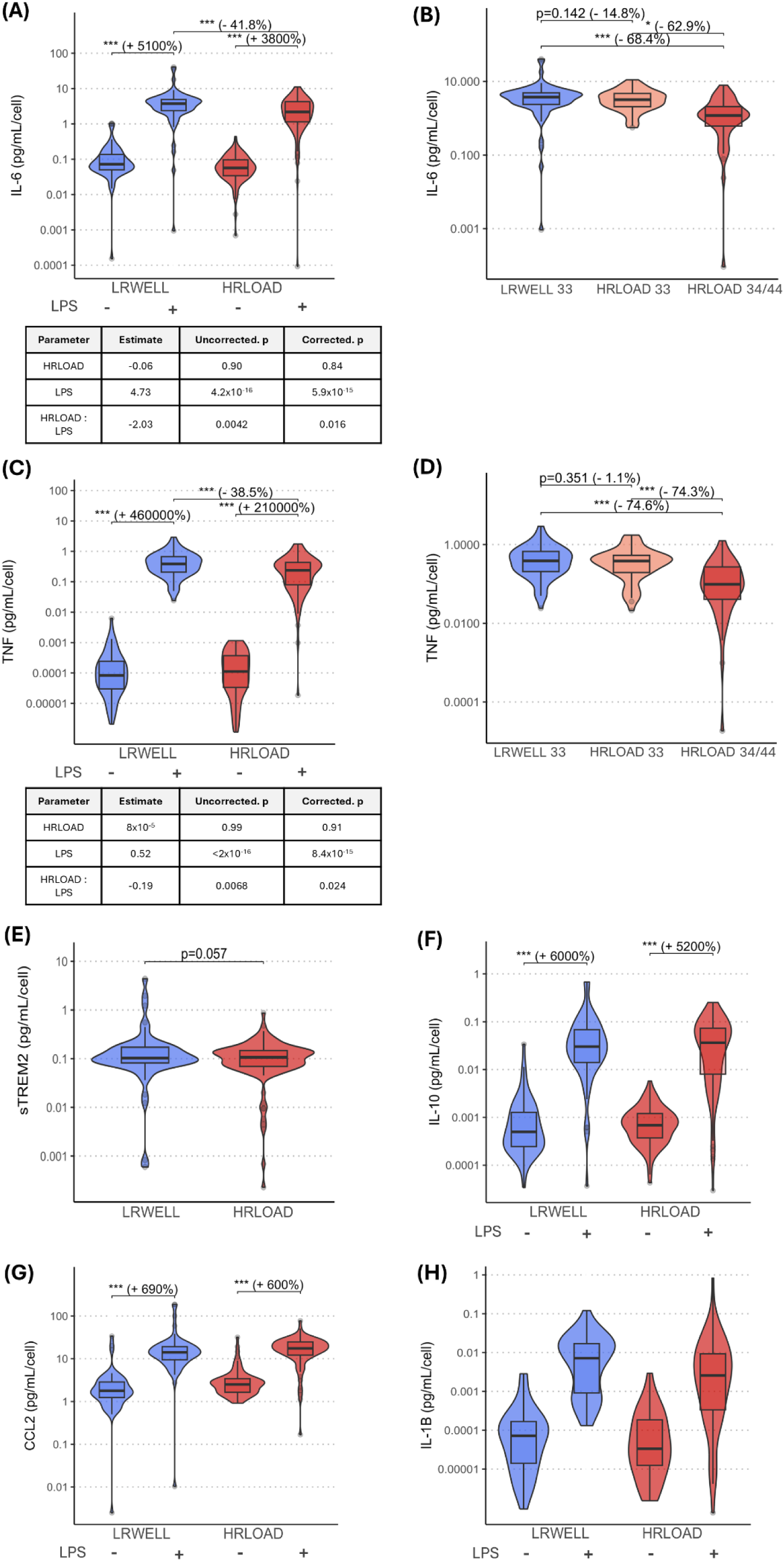
Impaired IL-6 and TNF inflammatory cytokine response in HRLOAD microglia driven by APOE ε4. Levels of secreted proteins were measured using a LegendPLEX cytometric bead array assay in unstimulated and LPS-challenged (20 ng/mL for 24 hours) iPSC microglia for (**A**) IL-6, **(C)** TNF, **(E)** sTREM2, (**F)** IL-10, **(G)** CCL2, **(H)** IL-1β. HRLOAD (red) was compared to LRWELL (blue), and LPS-challenged to unstimulated, using linear mixed effects modelling with p-values corrected by Benjamini-Hochberg procedure. Exploratory analysis of LPS-challenged data looking at the effect of APOE ε4 is shown for significant HRLOAD:LPS interactions in IL-6 (**B**) and TNF (**D**). LRWELL (blue), HRLOAD33 (orange), and HRLOAD34/44 (red) were compared using linear mixed effects modelling, with APOE as a fixed effect. Uncorrected p-values are displayed. Across all parts of the figure, data plotted is the mean of 4 replicate wells that were pooled, then normalized to the average cell count across those wells. Box plots display the median and interquartile range of secreted protein per cell (pg/mL/cell) for N = 28 HRLOAD, 15 LRWELL from a minimum of 4 experiments. * p < 0.05, ** p < 0.01, *** p < 0.001.

IBA1 is a widely used microglial marker whose expression increases in response to injury and is linked to phagocytic activity (Lier et al., 2021). HRLOAD microglia showed significantly lower IBA1 expression than LRWELL cells (uncorrected p= 0.02, corrected p= 0.05; Extended Data Figure 2), which was not significantly associated with *APOE ε4* status, despite most cells remaining IBA1-positive (Extended Data Figure 2). This reduction suggests an altered basal activation state in HRLOAD microglia.

### High polygenic risk iPSC microglia have a selective impairment in amyloid-β uptake

Endosomal genes are strongly implicated in genetic risk for AD (Maninger et al., 2024), and several late-onset AD risk genes are directly involved in phagocytosis. Assays measuring uptake of fluorescent cargo were used to assess endocytosis and phagocytosis. We observed significantly reduced endocytic uptake of soluble unaggregated amyloid-β in HRLOAD microglia (uncorrected p= 8 × 10^-4^, corrected p= 0.0047, −5.9%, Fig 3A), which was not dependent on *APOE ε4* (Extended Data Figure 3), and was partially inhibited by Dynasore, indicating some dynamin-mediated endocytic uptake (Fig 3B). In contrast, endocytic uptake of transferrin was not reduced and in fact trended towards an increase in HRLOAD microglia (uncorrected p= 0.025, corrected p= 0.058, Fig 3C). Phagocytic uptake of dead neuronal cells, myelin, and *E. coli* bioparticles were unaltered (Figs 3E, 3G and 3I). There were no differences in early (EEA1+) or late (Rab7+) endosome accumulation or early endosome size between the HRLOAD and LRWELL iPSC microglia (Extended Data Figure 3).

**Figure 3.**
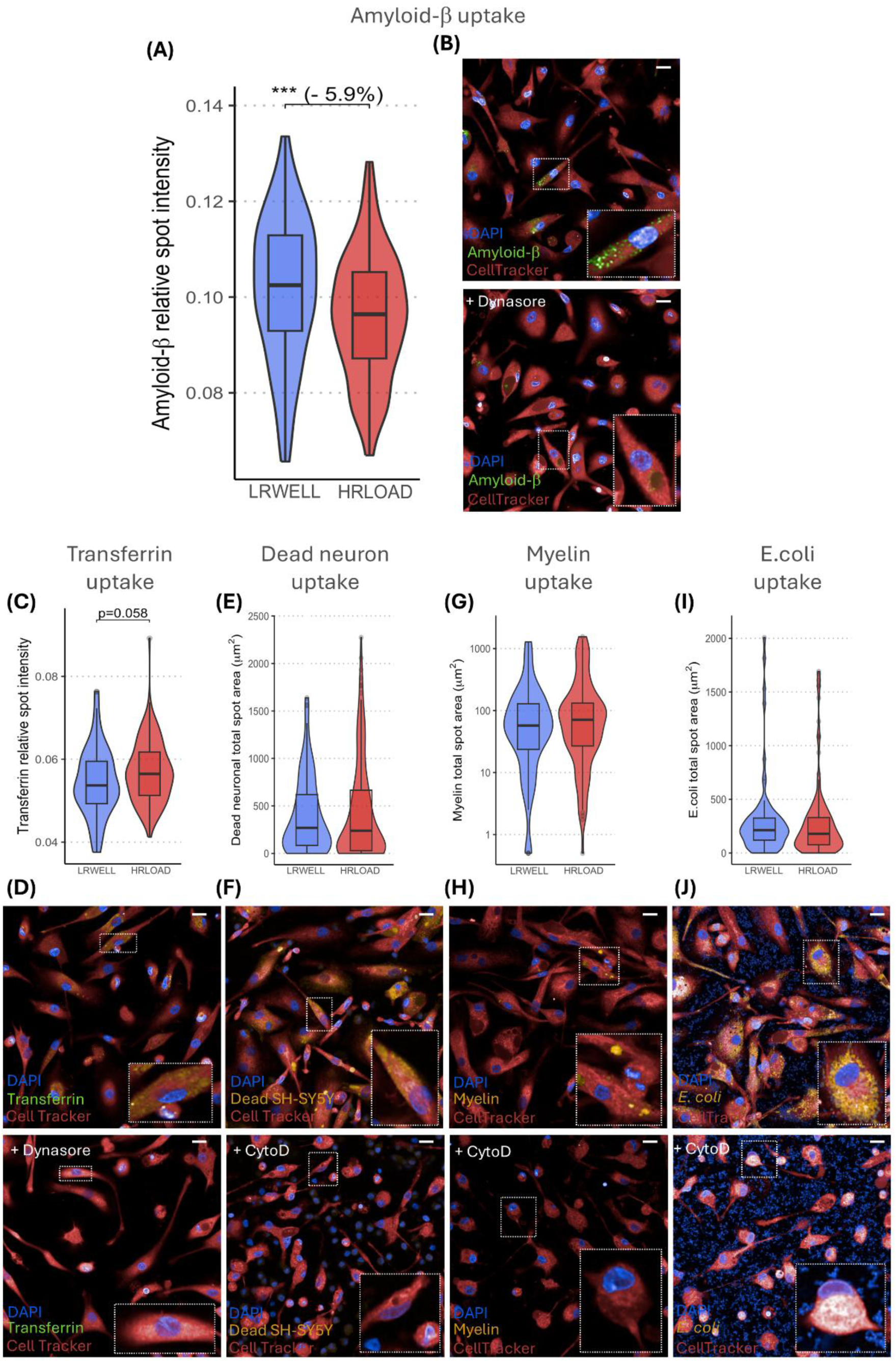
HRLOAD microglia have a selective defect in amyloid-β endocytic uptake. Endocytosis was investigated by performing uptake assays of **(A)** HiLyte-488 Amyloid-β 1-42 peptide and **(C)** pHrodo Red-transferrin, measuring relative spot intensity. Representative images (± Dynasore to inhibit endocytosis) are shown in **(B)** and **(D)** for each assay, respectively. Phagocytosis was investigated by performing uptake assays of pHrodo Red-labelled **(E)** dead neuronal cells, **(G)** mouse myelin, and **(I)** E. coli bioparticles, measuring total spot area per cell. Representative images (± Cytochalasin D to inhibit phagocytosis) are shown in **(F)**, **(H)** and **(J)** for each assay, respectively. Across all parts of the figure, a median of 4 technical well replicates was first taken in each experiment. Box plots display the median and interquartile range for N= 35 HRLOAD, 17 LRWELL from a minimum of 4 experiments. For all phenotypes HRLOAD was compared to LRWELL using linear mixed effects modelling with p-values corrected by Benjamini-Hochberg procedure. *** p < 0.001. Scale bars: 20 µm.

### Neutral lipid accumulation, calcium flux, and chemotaxis are not significantly altered in high polygenic risk iPSC microglia

Excessive neutral lipid accumulation is observed in AD plaque-associated microglia (Wu et al., 2025), so we assessed total cellular neutral lipid levels using HCS LipidTox Green Neutral Lipid Stain, using cyclosporin A as a positive control for lipid accumulation (Fig 4B). No significant difference was observed between HRLOAD and LRWELL microglia (Fig 4A). Purinergic signaling (via P2X/P2Y receptors) is disrupted in AD and is important for microglia inflammation, phagocytosis, and directed motility towards injured cells (Mei et al., 2024). To determine whether this pathway differs between LRWELL and HRLOAD iPSC-derived microglia, we quantified ATP-evoked intracellular calcium responses using Calcium-6 fluorescence imaging on the FLIPR platform (Fig 4D). Stimulation with ATP triggered a rapid rise in cytosolic calcium in both groups, with no significant differences observed between HRLOAD and LRWELL microglia (Fig 4C).

**Figure 4.**
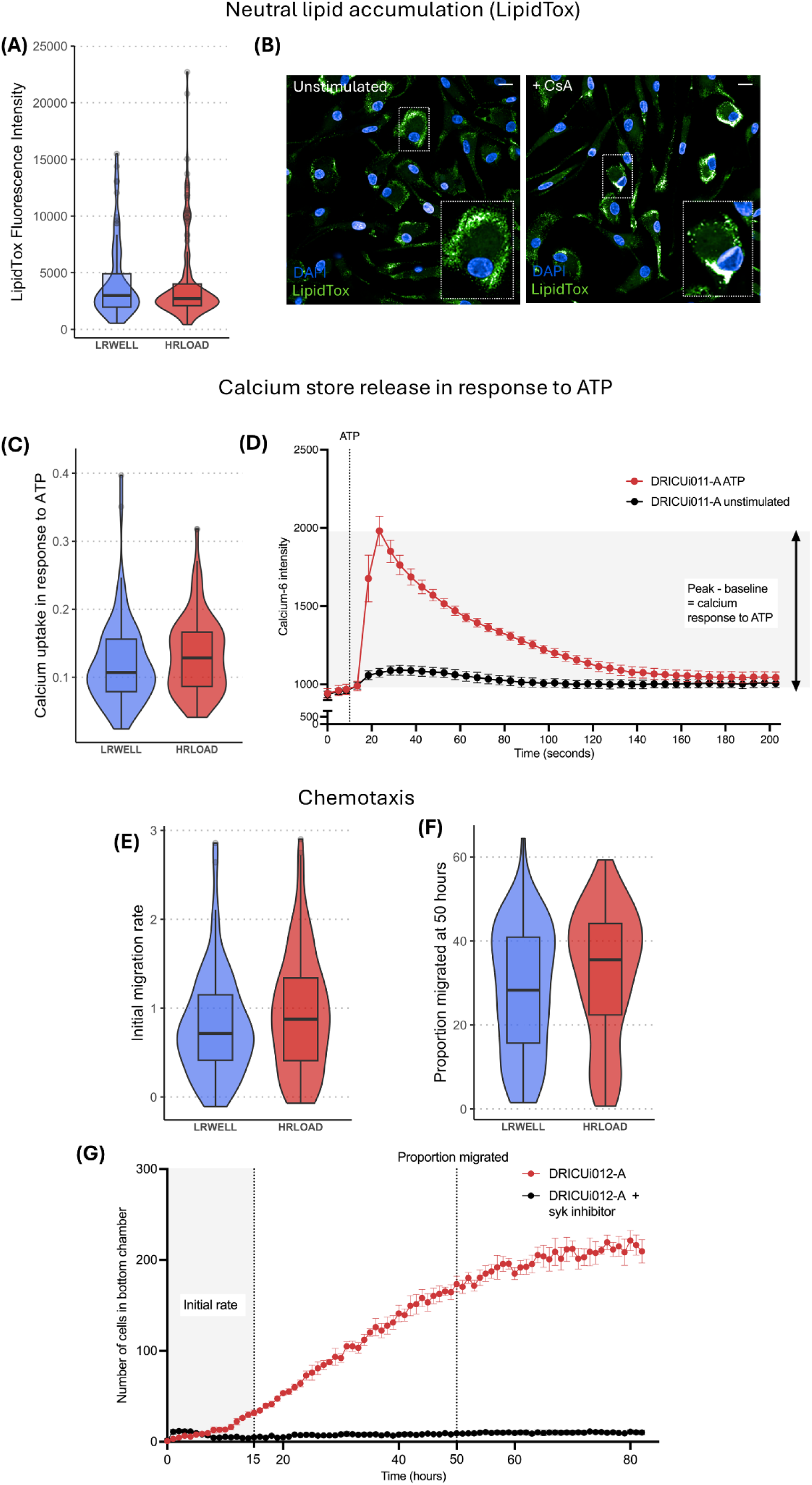
HRLOAD microglia do not exhibit alterations in neutral lipid accumulation, calcium flux, or chemotaxis. **(A)** Neutral lipids were investigated by staining microglia with HCS LipidTox Green, with mean fluorescence intensity per cell analysed. **(B)** Representative images of the LipidTox assay (± Cyclosporin A to induce neutral lipid accumulation). **(C)** Calcium flux was investigated by loading cells with Calcium-6 dye and stimulating with ATP, with the difference in peak versus baseline fluorescence analysed. **(D)** A representative trace of the calcium flux assay (± ATP). Data points show mean ± SEM of n=4 technical replicate wells from 1 assay. Chemotaxis was investigated using an IncuCyte chemotaxis assay with C5a as the chemoattractant, and the initial migration rate (proportion of total cells migrated per hour between 0-15 hours) (**E**) and proportion migrated at 50 hours (**F**) was analysed. **(G)** A representative trace of the chemotaxis assay (± Syk inhibitor to inhibit chemotaxis). Data points show mean ± SEM of n=4 technical replicate wells from 1 assay. Across all parts of the figure, a median of 4 technical well replicates was first taken in each experiment. Box plots display the median and interquartile range for N= 35 HRLOAD, 17 LRWELL from a minimum of 4 experiments. For all phenotypes HRLOAD was compared to LRWELL using linear mixed effects modelling with p-values corrected by Benjamini-Hochberg procedure. Scale bars: 20 µm.

Chemotaxis is an important activity of microglia that allows their directed migration towards pathology, for example amyloid plaques in the AD brain. Specific AD risk genes have been linked to microglia chemotaxis. We assayed microglial chemotaxis towards the chemoattractant C5a, using a Syk inhibitor as a negative control (Fig 4G). There was no significant difference between HRLOAD and LRWELL in the rate of chemotaxis or the maximum proportion of migrated cells (Figure 4E and 4F).

### Correlation analysis shows independence of metabolic, cytokine, and endocytosis functional effects of AD polygenic risk

Having identified high polygenic risk to have significant effects on immune challenged ATP production, challenged IL-6 and TNF secretion, basal IBA1 expression, and soluble amyloid-β uptake, we sought to test for relationships between these factors in the same lines using correlation analysis. We observed three independent functional clusters of correlation: (i) ‘metabolic’, consisting of challenged mitochondrial and glycolytic ATP production and basal IBA1 expression; (ii) ‘cytokine’, consisting of challenged IL-6 and TNF secretion; and (iii) ‘endocytosis’, consisting of amyloid-β and transferrin endocytic uptake (Fig 5A). This shows that polygenic risk influences these clusters independently. Within the metabolic cluster, challenged mitochondrial ATP production positively correlates with glycolytic ATP production (Fig 5B), but negatively correlates with basal IBA1 expression (Fig 5C). In the cytokine cluster, challenged IL-6 and TNF cytokine secretion show a positive correlation (Fig 5D). In the endocytosis cluster, amyloid-β and transferrin uptake are positively correlated (Fig 5E).

**Figure 5.**
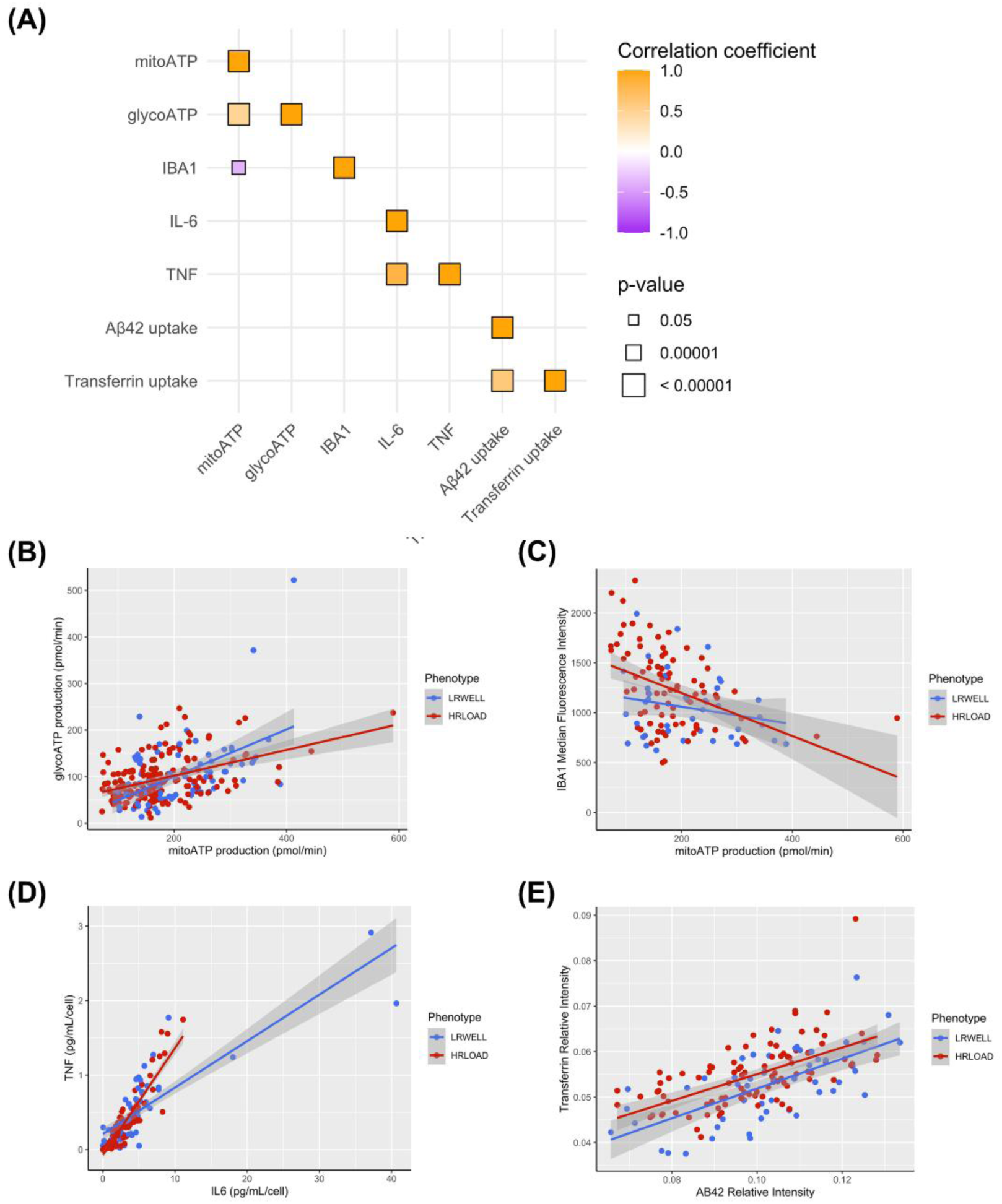
Correlation analysis reveals independent metabolic, cytokine, and endocytic functional effects. **(A)** Correlation matrix for the significant or trending HRLOAD phenotypes in this paper, which are LPS-challenged mitochondrial ATP production rate (mitoATP), LPS-challenged glycolytic ATP production rate (glycoATP), unstimulated IBA1 expression, LPS-challenged IL-6 secretion, LPS-challenged TNF secretion, amyloid-β (Aβ) uptake, transferrin uptake. P values include a Bonferroni correction for multiple testing. Individual correlation plots are shown for **(B)** mitoATP production versus glycoATP production, **(C)** mitoATP production versus IBA1 median fluorescence intensity, **(D)** IL-6 versus TNF secretion, **(E)** Aβ versus transferrin uptake. For (**B-E**), data points represent a median of n=3-10 technical well replicates from one iPSC line in an individual matched experiment.

### Proteomics of high polygenic risk microglia show significant changes in some biological pathways

Bulk proteomic analysis was performed on HRLOAD and LRWELL iPSC-microglia cell lysates with and without 24 hours LPS stimulation. Principal component analysis (PCA) shows clear separation in protein expression between the LPS-challenged and unstimulated samples along the first principal component, however HRLOAD and LRWELL overlap (Fig 6A). LPS stimulation alone caused a high number of differentially expressed proteins (Extended Data Figure 4), however when comparing HRLOAD to LRWELL there were no differentially-expressed proteins in the unstimulated samples, and only one differentially-expressed protein in the LPS-challenged samples, namely Proline Rich Protein BstNI Subfamily 1 (PRB1) (Fig 6B). We performed Gene Set Enrichment Analysis (GSEA) on all proteins to identify biological pathways which were significantly enriched following multiple testing correction. In unstimulated samples comparing high with low polygenic risk, the top 10 Reactome pathways ranked by log-fold change included significantly upregulated unfolded protein response (P= 0.002), cytokine (P=0.029), MAPK (P= 0.046) and GPCR signalling (P=0.028), and amino acid metabolism (P=0.018) (Fig 6C). No downregulated pathways were observed in the top 10. In LPS-challenged samples (Fig 6D) comparing high with low polygenic risk, the top 10 Reactome pathways by log-fold change included significantly upregulated rRNA modification (P=0.018) and processing (P=0.002), cell-cell communication (P=0.033) and calcium response (P=0.049), and significantly downregulated chromatin organization (P=0.003) and pyroptosis (P=0.049).

**Figure 6.**
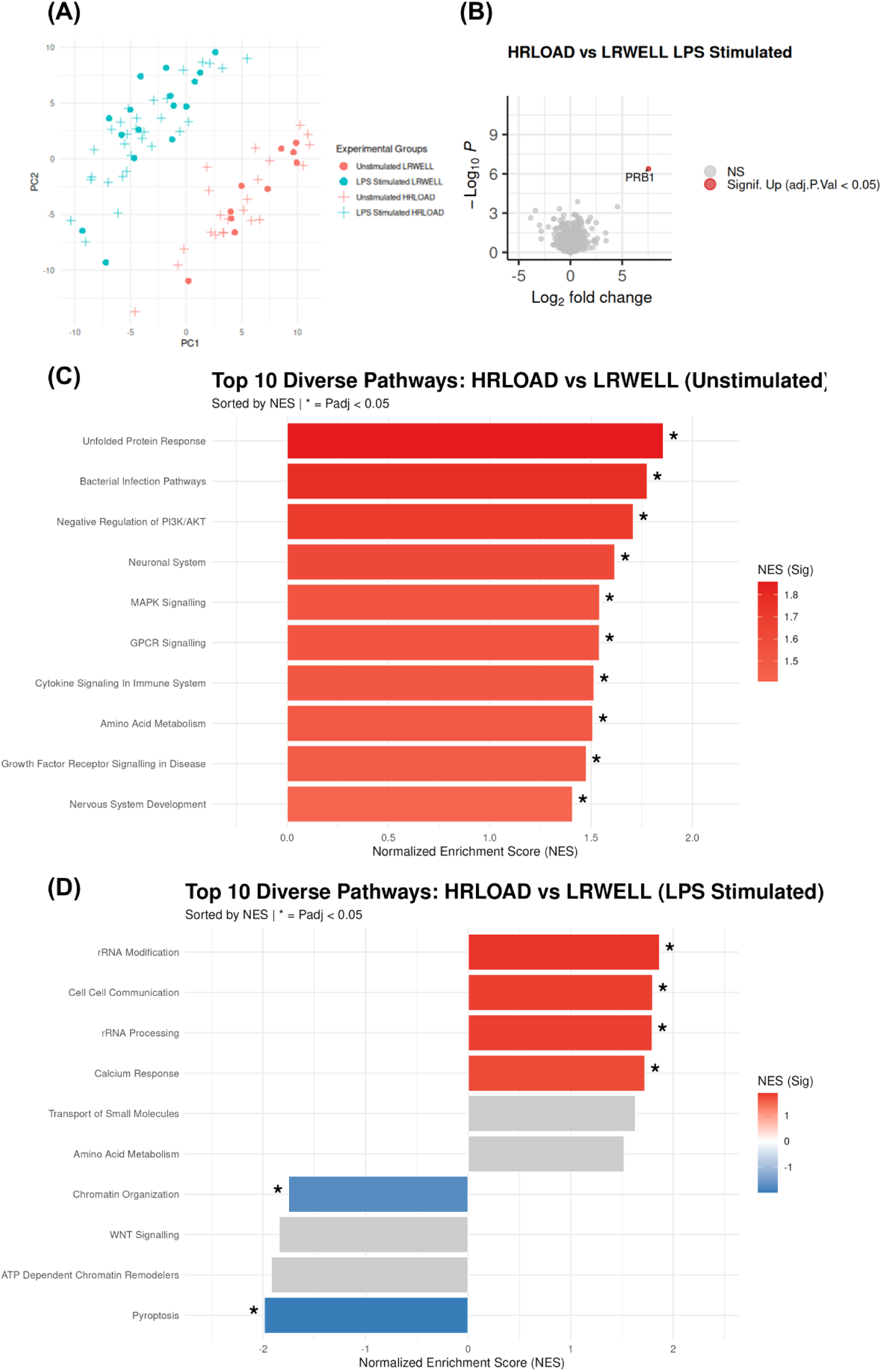
Proteomics analysis suggests dysregulated pathways in unstimulated and LPS-stimulated HRLOAD microglia. **(A)** Principal component analysis shows clear separation by stimulation condition but not by genotype. **(B)** Volcano plot of differentially expressed proteins comparing stimulated HRLOAD to LRWELL, with the significant protein highlighted in red. **(C,D)** Pathway Enrichment of the Proteome. Bar charts show the most statistically significant Reactome terms filtered by a Jaccard similarity coefficient (cutoff < 0.4) to eliminate redundant gene sets and highlight distinct biological processes for **(C)** Unstimulated and **(D)** LPS-stimulated (20 ng/mL for 24 hours) samples. Proteins were ranked for GSEA using a composite score of p-value and Log Fold-Change, * indicates significant enrichment, other terms are not statistically significant. N= 30 HRLOAD, 16 LRWELL. Data is from 2-3 technical replicate wells processed separately from one harvest.

Given the functional differences between HRLOAD and LRWELL cells, and the apparent independence of these effects from each other, we focused analysis on the proteomic signatures of microglia with extreme metabolic, cytokine or endocytic functional effects, to identify dysregulated biological pathways. We identified the bottom 5 HRLOAD and top 5 LRWELL microglia lines for (i) LPS-challenged mitochondrial ATP production (metabolic), (ii) LPS-challenged IL-6 secretion (cytokine), and (iii) amyloid-β uptake (endocytic), and performed GSEA comparison of HRLOAD to LRWELL for these extremes (Extended Data Figure 4). For the metabolic function there we observed a significant downregulation of respiratory complex I (P=0.001), mTORC1 activation (P=0.007), viral infection pathways (P=0.013), antigen processing (P=0.013), regulation of tRNA (P=0.003) and mRNA (P=0.013) and a significant upregulation of DNA-damage-induced senescence (P=0.013), antimicrobial peptides (P=0.007), metal ion response (P=0.004), and Rho GTPase (P=0.013) signalling. For the cytokine function, there was significant downregulation of cytokine signalling (P=0.001), particularly interleukin-1 (P=0.009) and pyroptosis (P=0.043) and decreased toll-like receptor activation (P=0.009), which mediate inflammatory signalling in response to detection of pathogen antigens such as LPS. Upregulated pathways were fewer and included class A1 GPCRs (P=0.049). For the endocytic function, we found a significant downregulation of cell cycle (P=1.6×10^-4^), telomere extension (P=0.001), mitochondrial translation (P=0.005), and ER stress (P=0.002) and a significant upregulation of small molecule/ion transport (P=0.006), glycosphingolipid catabolism (P=2.2x 10^-4^), reactive oxygen species production by phagocytes (P=0.001), and respiratory complex IV assembly (P=0.006).

We note that analysis of functional extremes must be treated as speculative and require replication in independent samples. Although lacking power to definitively identify dysregulated proteins, in this instance, proteomics can offer clues to underlying dysregulated cell processes that may be worthy of further exploration.

## Discussion

We tested the functional effects of iPSC microglia carrying a high burden of genetic risk for LOAD. We found microglia from high-risk individuals challenged by LPS stimulation exhibited significantly impaired mitochondrial ATP production and inflammatory cytokine release of IL-6 and TNF, with the latter driven by the presence of at least one *APOE ε4* allele. Under normal unstimulated conditions we observed a selective deficit in amyloid-β uptake. We also observed a small reduction in IBA1 expression, suggesting alteration to the basal activation state of the microglia. Correlation analysis performed on the significant main effects of high polygenic risk for AD showed independence of three functions namely, ‘metabolic’, ‘cytokine’ and ‘endocytic’, which are summarized graphically in Figure 7. Previous studies of AD polygenic risk in iPSC have shown associations with reduced interferon response protein expression in astrocytes (Lee et al., 2025), and a correlation with MHC class II gene expression in microglia xenotransplanted into *App^NLGF^* mice (De Strooper et al., 2025).

**Figure 7.**
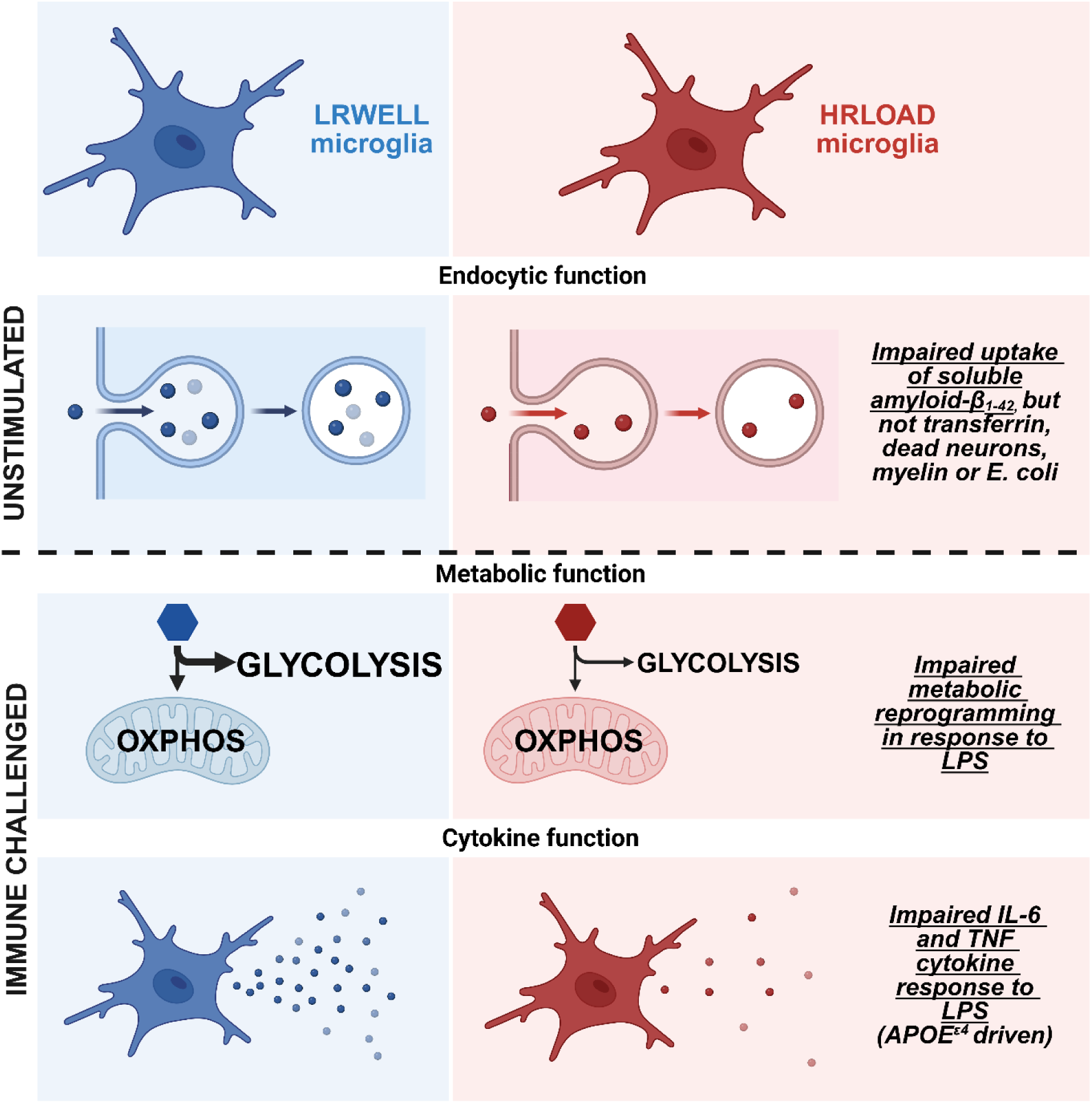
Effect of AD high polygenic risk on iPSC microglia. Summary graphical illustration of main HRLOAD microglia functional phenotypes reported in this paper. Created in BioRender. O’Donoghue, R. (2026) https://BioRender.com/0wvs2go

The impairment of metabolic function in HRLOAD microglia following immune challenge is characterized by reduced mitochondrial and glycolytic ATP production. Microglia mainly produce ATP through mitochondrial oxidative phosphorylation at rest but switch to upregulate glycolytic non-mitochondrial ATP production when activated by a stimulus, a process known as “metabolic reprogramming” or “metabolic switching” (Miao et al., 2023). This metabolic flexibility allows microglia to respond quickly to stimuli, as glycolysis is faster although less efficient, than mitochondrial production of ATP. These findings may reflect an underlying mitochondrial defect. MMP of AD microglia was slightly impaired at baseline, which suggests alterations to mitochondrial respiration at rest (Connolly et al., 2018). Furthermore, proteomics analysis suggested that microglia with the ‘metabolic’ deficit may have impaired complex I biogenesis and activity of mTOR, which is a positive regulator of mitochondrial biogenesis and respiration. Supporting this, a recent pre-print that included 14 of the iPSC lines from this study found an association between high AD PRS and reduced expression of oxidative phosphorylation genes (Perez-Alcantara et al., 2025). Mitochondrial impairments have been observed in neuronal cybrid models containing late-onset AD mitochondria, which showed reduced MMP and respiration rates (Samanta et al., 2023; Trimmer et al., 2000). Furthermore, a rare AD risk variant of *TREM2* has been linked to impaired metabolic reprogramming in iPSC-microglia (Piers et al., 2019). Also, in a proteomic study of over 2,000 human brains, the protein expression module most strongly associated with AD progression implicated microglia and astrocyte sugar metabolism and was enriched in AD genetic risk variants (Johnson et al., 2020). Surprisingly, correlation analysis revealed that mitochondrial ATP production had a weak but significant negative correlation with basal IBA1 expression. IBA1 expression is a biomarker of activated microglia and is hypothesised to have a role in phagocytosis (Kenkhuis et al., 2022; Lier et al., 2021). The mechanism is unclear, particularly as IBA1 was measured in the absence of LPS stimulation, whereas mitochondrial ATP production was altered under LPS stimulation.

A distinct ‘cytokine’ component was identified in HRLOAD microglia, consisting of blunted pro-inflammatory cytokine responses to LPS stimulation, which were positively correlated. It is tempting to speculate a connection between this and the impaired metabolic reprogramming effect, especially as prior literature shows that glycolysis controls microglia activation and cytokine production (Codocedo et al., 2024; Pålsson-McDermott & O’Neill, 2020). However, we did not observe a significant correlation between ATP production rates and inflammatory cytokine secretion, suggesting that these are independent. We also found that *APOE ε4* is the driver of the blunted pro-inflammatory cytokine responses to HRLOAD, but is not significantly associated with impaired metabolic reprogramming. further supporting the independence of these two effects. It seems paradoxical that high genetic risk for AD reduces activation and pro-inflammatory response, as neuroinflammation is believed to be a major driver of disease progression, however our findings align with previous studies of the AD-associated risk variant R47H in *TREM2*, which have shown that R47H TREM2 impairs LPS-stimulated metabolic reprogramming and microglia activation, reduces microglia reactivity at amyloid plaques and plaque compaction, and increases axonal dystrophy around amyloid deposits (Piers et al., 2019; Song et al., 2018; Yuan et al., 2016). Therefore, human genetic evidence combined with *in vitro* and *in vivo* modelling suggests that reduced microglia activation is detrimental in AD.

Amyloid-β uptake was significantly impaired in HRLOAD microglia. As part of a distinct ‘endocytic’ function, this has a strong positive correlation with transferrin uptake, yet transferrin uptake was not impaired, indeed it showed a moderate increase that was not significant after multiple testing correction. Our interpretation is that HRLOAD microglia do not have a universal impairment to endocytic uptake, instead there is a selective deficit in amyloid-β uptake associated with AD polygenic risk. This may suggest a reduction in cell-surface receptors that mediate amyloid-β recognition, such as TREM2, SR-A, MER and AXL (Fu et al., 2025). Amyloid-β uptake by microglia is important for the formation of dense-core amyloid plaques, which are proposed to be a mechanism for confining and restricting amyloid-β to limit the spreading of neurotoxic amyloid-β oligomers (Huang et al., 2021). Therefore, the finding that human microglia with high polygenic risk for AD have reduced amyloid-β uptake suggests they have poorer ability to both clear amyloid-β and construct dense-core amyloid plaques, thereby providing a clear and primary mechanism underpinning AD development.

Proteomics analysis added further clues to the nature of convergent dysregulated protein expression in the HRLOAD microglia. Only one significant differentially-expressed protein was identified-PRB1, a poorly-characterized salivary protein that was upregulated. When we explored the most dysregulated biological pathways via gene set enrichment analysis we observed HRLOAD under immune challenge was associated with significantly downregulated pyroptosis. Pyroptosis is a type of programmed cell death that is highly inflammatory, involving inflammasome-mediated IL-1β production, therefore downregulated pyroptosis is consistent with reduced inflammatory activation (Bergsbaken et al., 2009). Although IL-1β secretion was not significantly reduced by high polygenic risk in cytokine assays, further exploration with additional techniques may be necessary to validate potential alterations to pyroptosis. Furthermore, unstimulated HRLOAD microglia had significantly upregulated unfolded protein response and amino acid metabolism, alongside perturbed cytokine signaling. The unfolded protein response causes changes to amino acid metabolism and global translation, and sustained activation can lead to alterations in inflammatory signalling, autophagy, mitochondrial function and programmed cell death pathways (Gawlak-Socka et al., 2026; Grootjans et al., 2016). Therefore, the unfolded protein response is an interesting target for future follow-up, and this provides further support for the hypothesis that high polygenic risk alters the baseline activation state of microglia.

Limitations should be considered when interpreting these findings. iPSC-derived microglia generated in monoculture resemble a more fetal-like (Abud et al., 2017), partially activated state (Cadiz et al., 2022) rather than a fully mature adult profile, reflecting the absence of cues from surrounding brain cells. *In vivo*, microglia interact continuously with neurons, astrocytes, oligodendrocytes, and vasculature, each of which would themselves carry high or low AD polygenic risk. Nevertheless, isolating microglia provides a valuable window into the cell-autonomous consequences of AD genetic risk, particularly given the strong enrichment of risk genes in microglial pathways. A further limitation is that all donor lines were derived from white European ancestry (Maguire et al., 2025).

We show for the first time in a human *in vitro* model of LOAD, that high polygenic risk under immune challenge causes weaker metabolic reprogramming, diminished inflammatory responses (*APOE ε4* driven), and under basal conditions, a selective impairment of amyloid-β uptake. We argue that these microglial deficits may represent primary events in the pathogenesis of LOAD. Indeed, the reduced uptake of amyloid-β supports that these deficits in microglia predate, and contribute to, plaque formation. Thus, intervention at this early stage to modify microglial deficits must be a major focus for future therapeutics. Our findings also validate this approach as an *in vitro* human iPSC platform to test drugs and genetic therapies for the most common form of Alzheimer’s disease.

## Methods

### Polygenic Risk Selection

Details of genotyped samples and QC were performed as detailed in (Maguire et al., 2025). PRS calculation methods are described in (Maguire et al., 2025) and also detailed in the Supplementary Methods. (Leonenko et al., 2021). AD cases (HRLOAD) were selected on an age of onset threshold >65, and with a high PRS (Mean PRS_HRLOAD_ = 2.21, SD PRS<u>_HRLOAD_</u> = 0.48,, Max PRS<u>_HRLOAD_</u> = 3.23, Min<u>_HRLOAD_</u> = 1.25), while control individuals (LRWELL) were selected if they had an APOE ε3/3, age at interview > 70 y.o., (Mean age at interview = 80.7 y.o., Min_AGE_ = 71, Max_AGE_ = 95), and with a low LOAD PRS (Mean PRS_LRWELL_ = −1.8, SD PRS<u>_LRWELL_</u> = 0.5,, Max PRS<u>_LRWELL_</u> = −1.03, Min<u>_LRWELL_</u> = −2.83). We selected a rough balance of males and females.

### Induced pluripotent stem cell lines

The power of this study is based upon sampling from the extremes of polygenic risk for AD and the use of multiple donors at each extreme. It is appreciated that individuals will have different genetic backgrounds in addition to sharing AD genetic risk and that stem cell research is particularly subject to the effects of variation (Volpato et al., 2018). Thus, a powerful study is essential to differentiate disease effects from background variation. The iPSC lines used derived from peripheral blood mononuclear cells (PBMC) using non-integrating Sendai reprogramming vectors (Cytotune, Life Technologies), from donors in the Alzheimer’s Disease Cardiff Cohort comprising 5,000 well-characterised AD cases and 3,000 cognitively well aged controls (Maguire et al., 2025). A total of 51 iPSC lines were used: 34 LOAD cases and 17 cognitively well, aged controls. Male (46% of total) and female (54% of total). donors were selected based on their global AD PRS and age of onset, with the LOAD-diagnosed donors having an age of onset over 65 and a high global AD PRS (mainly >1.8 SD), and the cognitively well donors having an age at participation over 70 and a low global AD PRS (mainly <-1.8 SD). All cognitively well donors were *APOE* ε3/3, whereas the LOAD donors were 53% *APOE* ε3/3, 32% *APOE* ε3/4, 15% *APOE* ε4/4.The PBMC were collected with informed consent under four studies with REC IDs 12/WA/0052, 04/9/030, 17/SS/0139, 00/09/42, and the present study was approved by the research ethics committee Wales REC 3 (REC ID 12/WA/0052). iPSC-microglia were differentiated from 6 batches of iPSC lines, with three “batch controls” present in every differentiation, using a previously published protocol with minor modifications (Washer et al., 2022), which is detailed in the Supplementary Methods. The iPSC lines are listed in Supplementary Table 1, with more details of their characterisation in (Maguire et al., 2025) and in hPSCreg, https://hpscreg.eu/. Most lines were deposited in the European Bank for Induced Pluripotent Stem Cells, EBiSC. iPSCs were maintained in mTeSR-Plus (STEMCELL Technologies) on hESC-qualified Geltrex-coated (Gibco) plates and passaged as clumps with 0.5 mM EDTA (Sigma) in PBS.

### Proteomics

iPSC-microglia were differentiated for 8 days (300,000 cells/well in 6 wells of 12-well plate), with a half-media change on day 7. On day 8, a full media change was performed with 1 mL of full Microglia media +/- *E. coli* lipopolysaccharide (20 ng/mL, Invivogen, tlrl-eklps). Following 24 hours incubation, cells were washed twice with PBS, then lysed in 400 µL lysis buffer (5% SDS, 10 mM Tris-(2-Carboxyethyl)phosphine (TCEP), 50 mM triethylammonium bicarbonate (TEAB) buffer), scraped into 1.5 mL low-binding tubes, and stored at −80 °C. Mass spectrometry was performed as described in the Supplementary Methods.

### Immunocytochemistry

iPSC-microglia were plated at a density of 10,000 cells/well in fibronectin coated ½ area 96-well plates. A half media change was performed on day 7. On day 10, media was removed and iPSC-microglia were fixed with 4% paraformaldehyde (Thermo Scientific) for 10 minutes at room temperature. Fixed cells were washed 3 times with PBS, and stored at 4°C until immunocytochemistry was performed. Two panels of immunocytochemistry were carried out for these cells: microglia and endocytosis. For immunocytochemistry, blocking buffer of 10% donkey serum (Abcam) in 0.1% (microglia) or 0.4% (endocytosis) Tween-20 (VWR) in PBS was added for 1 hour at room temperature. Following this, block was removed and replaced with primary antibodies in the panel specific blocking buffers. For microglia, IBA1 (1:100) was used. For endocytosis, IBA1 (1:500), EEA1 (1:500) and Rab7 (1:500) were used. Primary antibodies were incubated at 4°C overnight. The following day, primary antibodies were removed and cells washed 3 times with 0.1% (microglia) or 0.4% (endocytosis) Tween-20 in PBS. Secondary antibodies and DAPI (5 µg/mL), diluted in the panel-specific Tween-20 concentration in PBS, were then added and incubated for 1 (microglia) or 2 (endocytosis) hours at room temperature. For microglia, Alexa Fluor-488 donkey anti-goat was used at 1:500 dilution. For endocytosis, Alexa Fluor-488 donkey anti-goat (1:1000), Alexa-Fluor-568 donkey anti-rabbit (1:1000) and Alexa-Fluor-647 donkey anti-mouse (1:1000) were used. Secondary antibodies were washed off in the same way as primary antibodies. Finally, PBS was added to the wells, and single z-plane imaged (microglia: 40x water objective, confocal, 0 µm, 15-17 fields per well; endocytosis: 63x water objective, confocal, 0 µm, 9 fields per well) using an Opera Phenix. Images were analysed using the Columbus software. For microglia, percentage of cells stained positive for IBA1 and the mean cellular intensity of IBA1 were calculated. For endocytosis, spots of EEA1 and RAB7 were segmented and the area and intensity of spots were calculated.

| ANTIBODIES AND STAINS | SOURCE | IDENTIFIER |
| --- | --- | --- |
| IBA1 | Abcam | ab5076 |
| EEA1 | Abcam | ab70521 |
| Rab7 | Abcam | ab137029 |
| Alexa Fluor-488 Phalloidin | Life Technologies | A12379 |
| Donkey anti-goat 488 | Life Technologies | A11055 |
| Donkey anti-rabbit 568 | Life Technologies | A10042 |
| Donkey anti-mouse 647 | Life Technologies | A31571 |

### Seahorse XF Real Time ATP rate assay

For Seahorse metabolic flux assays, oxygen consumption rates (OCR) and extracellular acidification rates (ECAR) were measured with the XF96 Extracellular Flux Analyzer, using the Seahorse XF Real-Time ATP rate assay kit, according to the manufacturer’s instructions (Agilent Technologies). iPSC-microglia were differentiated for 8 days (300,000 cells/well in 2-4 wells of a 12 well plate). On day 8, cells were lifted by incubating with TrypLE Express for 8 minutes at 37°C, 5% CO_2_, collected in a tube, centrifuged at 400×g for 5 minutes, resuspended in Microglia media, and counted. Cells were replated at 40,000 cells/well in fibronectin-coated (10 µg/mL, 30 minute minimum coating time) XFe96 Pro Cell Culture Microplates, using 3-10 wells per iPSC line in duplicate plates. Later on the same day, one of the duplicate plates received LPS treatment (20 ng/mL, 18 hours). On day 9, microglia media was removed from XFe96 Pro Cell Culture Microplates and replaced with 100 µL/well Seahorse XF medium (non-buffered Dulbecco’s modified Eagle’s medium containing 2.5 mM glucose, 2 mM L-glutamine, and 1 mM sodium pyruvate), prior to 45 minute incubation in a zero CO_2_ 37°C incubator. During this incubation, chemicals required for the ATP rate assay were prepared and loaded into XFe96 Pro sensor cartridges according to manufacturer’s instructions, prior to placing the sensor cartridge into a zero CO_2_ 37°C incubator. After the 45 minute incubation, the XFe96 Pro Cell Culture Microplates containing microglia in Seahorse XF medium were removed from the zero CO_2_ 37°C incubator and imaged and analysed for whole-well cell counts using the Celigo Image Cytometer (Revvity). Following imaging, all media was removed from the XFe96 Pro Cell Culture Microplates containing microglia and replaced with 180 µL/well of fresh Seahorse XF medium, prior to placing into a zero CO_2_ 37°C incubator. Both XFe96 Pro Cell Culture Microplates and drug loaded XFe96 Pro sensor cartridges were then inserted into the Seahorse XF96 analyser according to manufacturers instructions, and OCR and ECAR readings were acquired both at baseline and following addition of oligomycin and rotenone/antimycin A. For LPS-treated cells, experiments were timed so that baseline OCR and ECAR readings were taken by the XF96 Extracellular Flux Analyzer following 18 hour stimulation.

### Mitochondrial membrane potential and morphology assay

Mitochondrial morphology measured using Mitotracker Green (Invitrogen, M7514) and mitochondrial membrane potential was measured using JC-1 (Invitrogen, T3168). iPSC-microglia were differentiated for 10 days (7000 cells/well in 8 wells of a PhenoPlate 384-well microplate (Revvity)). On day 10, the cell media was removed and 4 wells were stained with 50 µL of a mixture NucBlue (1 drop/mL, Molecular Probes, R37605) and Mitotracker Green (Fisher M46750, stock 1mM in DMSO, final concentration 200 nM) in Live Cell Imaging Solution (Invitrogen, A59688DJ) at 37°C for 30 minutes. The stain was removed and replaced with 50 µL of Live Cell Imaging Solution, then z-stacks (40x confocal, 5 planes, 1 µm apart) imaged in an Opera Phenix at 37°C, 5% CO_2_. Later on day 10, 4 wells (not previously stained) were stained with 50 µL of a mixture NucBlue (1 drop/mL) and JC-1 (5 µg/mL) in Live Cell Imaging Solution at 37°C for 20 minutes. FCCP (1 µM, Cayman Chemical, CAY15218) was added as a positive control to spare wells alongside the staining mixture. The stain was removed and replaced with 50 µL of Live Cell Imaging Solution, then a single z-plane imaged (40x confocal, 0 µm, 15 fields per well) in an Opera Phenix at 37°C, 5% CO_2_. Automated image analysis was performed in Columbus software. For an estimation of mitochondrial mass, a maximum projection of the Mitotracker Green images was analysed to determine the average fluorescence intensity of the Mitotracker Green per cell. For an estimation of the mitochondrial network width:length ratio, an SER Ridge filter was applied to the maximum projection of the Mitotracker Green images and used to segment the mitochondrial network in each cell, before calculation of the ratio of width to length. To quantify the mitochondrial membrane potential, the average fluorescence intensity of the green and red fluorescence signals per cell were separately determined, then the ratio of red:green fluorescence intensity was calculated, where red fluorescence represents JC-1 aggregates in healthy polarized mitochondria and green fluorescence is monomeric JC-1 in the cytoplasm.

### LegendPLEX cytometric bead array assay

iPSC-microglia were differentiated for 9 days (20,000 cells/well in 4 wells of a 96-well microplate (Corning)). On day 9, a full media change was performed with 150 µL of full microglia media +/- *E. coli* lipopolysaccharide (20 ng/mL, Invivogen, tlrl-eklps). Following a 24 hour incubation, the cell supernatants were collected, combining the 4 replicate wells into a single tube which was stored at −80 °C. The microglia were then fixed by incubation with 4% paraformaldehyde for 10 minutes at room temperature and washed three times with 100 µL PBS. Cell nuclei were stained with DAPI (5 µg/mL in PBS) for 10 minutes at room temperature, washed twice with PBS, and then imaged and analysed for whole-well cell counts using the Celigo Image Cytometer (Revvity).

Cell supernatants were analysed for the concentration of secreted IL-6, TNF, IL-1β, IL-10, CCL2 (MCP-1), and sTREM2 in duplicate using a custom human panel LegendPLEX assay (Biolegend) and an Attune NxT flow cytometer with CytKick Autosampler, according to the manufacturer’s instructions and using LEGENDplex software (Biolegend) for the data analysis. For each analyte of each sample, a mean of the technical replicates was calculated, and normalized by division with the average cell count (average of 4 replicate wells).

### LipidTOX assay

Neutral lipid accumulation was measured using HCS LipidTOX Green (Invitrogen, H34475). iPSC-microglia were differentiated for 9 days (7000 cells/well in 4 wells of a PhenoPlate 384-well microplate (Revvity)). On day 9, the cell media was removed and the cells fixed by incubation with 50 µL of 4% paraformaldehyde for 10 minutes at room temperature. The paraformaldehyde was removed and the cells washed once with PBS, then stained with 50 µL of a mixture of NucBlue (1 drop/mL) and HCS LipidTOX Green (1:1000) in PBS at 37°C for 30 minutes. The plate was imaged on a single z-plane without washing in an Opera Phenix (40x confocal, 1 µm, 19 fields per well). Automated image analysis was performed in Columbus software to determine the average fluorescence intensity of the LipidTOX Green per cell.

### Calcium assay

Calcium release was measured using the FLIPR Calcium 6 assay (Molecular Devices, R8190). iPSC-microglia were differentiated for 9 days (7000 cells/well in 4 wells of a PhenoPlate 384-well microplate (Revvity)). A 2X working solution of Calcium 6 was prepared according to manufacturer’s instructions. On day 9, the cell media was removed and 8 wells were stained with 50 µL of Calcium-6 dye diluted 1:1 in Live Cell Imaging Solution at 37°C for 2 hours. A solution of 150 µg/mL ATP was prepared in Live Cell Imaging Solution and 50 µL added to 4 well of an empty 384-well source plate; Live Cell Imaging Solution alone was added to a further 4 wells as a no-stimuli control. Both plates were loaded onto a FLIPR Penta (Molecular Devices), and the plate containing iPSC-microglia was imaged with excitation 470-495 nm and emission 515-575 nm to establish a baseline reading (gain=80, exposure time= 0.15s, excitation intensity= 50%). Next, 25 µL of either the ATP solution or the Live Cell Imaging Solution control was removed from the source plate and dispensed into the corresponding wells of the imaging plate at a height of 50 µL, speed 25 µL/s (pipettor held during dispense, removal speed 20 mm/s). This resulted in a final concentration of ATP in the experimental wells of the iPSC-microglia plate of 50 µg/mL. The FLIPR imaged once per second, 10 times before ATP addition and 200 times after addition. The raw data was exported to MS Excel and the baseline signal (prior to ATP addition) subtracted from the peak calcium signal.

### Chemotaxis assay

Chemotaxis was measured using the IncuCyte Chemotaxis assay (Sartorius). iPSC-microglia were differentiated for 10 days (300,000 cells/well in 1 well of a 12-well plate), with a half-media change on day 7. On day 10, cells were lifted by incubating with StemPro Accutase for 15 minutes at 37°C, 5% CO_2_, collected in a tube, centrifuged at 400×g for 5 minutes, resuspended in Microglia media, and counted. Cells were replated at 2000 cells/well in 60 µL, in a fibronectin-coated (10 µg/mL, 30 minute minimum coating time) IncuCyte ClearView 96-Well Cell Migration Plate, using 3 wells per iPSC line, with the Syk inhibitor BIIB-057 (5 µM, Cayman, 17652) used in a few spare wells as a positive control. After allowing 30 minutes for the cells to settle at room temperature, Microglia media containing chemoattractant (1 nM recombinant human C5a, Peprotech, 300-70) was added to all wells of the reservoir plate underneath the transwell insert, 200 µL/well. The plate was loaded into an IncuCyte S3 Live-Cell Analysis System (Sartorius), and imaged using the Chemotaxis software module (10X, phase) with repeated scans every 1 hour for 4 days. The proportion of migrated cells for each cell line in each experimental replicate was calculated as the number of cells in the bottom half of the transwell over the total number of cells in both top and bottom compartments multiplied by 100. The initial rate of migration was calculated by finding the difference in proportion of cells at 0 hours and 15 hours divided by the 15 hours passed (i.e. proportion of total cells migrated per hour). Proportion of cells migrated at 50 hours was also used as an analysis metric as the approximate point of plateau in migration.

### Endocytosis assays

#### Cargo preparation

Endocytosis of two different substrates was analysed: pHrodo Red Transferrin (Invitrogen, P35376) and Beta-Amyloid (1-42), HiLyte™ Fluor 488-labeled (Anaspec, AS-60479-01). The pHrodo Red Transferrin was reconstituted in deionized water to 5 mg/mL. Stocks were stored at 4°C. The HiLyte-488 beta-amyloid was reconstituted in 50 µL 1% NH_4_OH, then diluted to 0.1 mM in PBS. Stocks were aliquoted and stored at −20°C.

#### Assay

iPSC-microglia were differentiated for 10 days (7000 cells/well in 12 wells of PhenoPlate 384-well microplate (Revvity)). On day 10, the iPSC-microglia were stained with 30 µL of a mixture of NucBlue (1 drop/mL) and CellTracker Deep Red plasma membrane stain (2 µM, Invitrogen, C34565) in Live Cell Imaging Solution (Invitrogen) with 20 mM Glucose (LCIS^+^) for 30 minutes at 37°C, 5% CO_2_. Cells were washed once with 50 µL of LCIS^+^, and then 50 µL of LCIS^+^, with the dynamin inhibitor Dynasore (30 µM for 30 minutes, Tocris, 1233) used in a few wells as a positive control. To initiate endocytosis, 25 µL of each endocytic substrate was added to 4 wells per iPSC line, so that a final well concentration of 25 µg/mL pHrodo Red Transferrin or 0.25 µM HiLyte-488 beta-amyloid. The plates were incubated for 3 hours at 37°C, 5% CO_2_, and a single z-plane imaged (40x water objective, confocal, 1 µm, 9 fields per well) in an Opera Phenix at 37°C, 5% CO_2_. Automated image analysis pipelines in Columbus software were used to segment spots of endocytic cargo within the iPSC-microglia (CellTracker and NucBlue used to identify and segment individual microglia), with medians per well obtained for the relative spot intensity, a parameter chosen because it best reflects the quantity of soluble endocytic cargo in endosomes.

### Phagocytosis assays

#### Cargo preparation

Phagocytosis of three different substrates was analysed: pHrodo Red E coli bioparticles (Invitrogen, P35361), paraformaldehyde-fixed SH-SY5Ys labelled with pHrodo iFL Red STP Ester (Invitrogen, P36011), and mouse brain myelin labelled with pHrodo iFL Red STP Ester (Invitrogen). The SH-SY5Ys were prepared fresh for each individual experiment as described in (Hall-Roberts et al., 2021), in brief: live SH-SY5Ys were dissociated, washed with PBS, fixed in 2% paraformaldehyde (4% paraformaldehyde diluted 1:1 with Live Cell Imaging Solution) for 10 minutes at room temperature, washed once with Hanks Buffered Saline Solution (Invitrogen), labelled with 40 µg/mL pHrodo iFL Red STP Ester (P36011, Invitrogen) for 15 minutes at room temperature, and washed twice before resuspending in Microglia media (400,000 cells/mL). The mouse brain myelin was extracted from brains of 9 month old female App-KI mice using the sucrose gradient fractionation protocol described in (Andreone et al., 2020), adjusted to 1 mg/mL protein concentration using the Protein Assay Kit II (BioRad, 5000112), and aliquots frozen at −80°C in HEPES-buffered media with 5% (v/v) DMSO. Prior to the first experiment of a differentiation, a 100 µg aliquot of mouse brain myelin was thawed, centrifuged at 4000×g for 10 minutes, resuspended at 1 mg/mL in Live Cell Imaging Solution (Invitrogen), labelled with 40 µg/mL pHrodo iFL Red STP Ester (P36011, Invitrogen) for 15 minutes at room temperature, and washed twice before resuspending in Microglia media (0.77 mg/mL). Labelled myelin was stored at 4°C for up to 4 weeks and the same stock used for the experimental repeats within a differentiation.

#### Assay

iPSC-microglia were differentiated for 9 days (7000 cells/well in 12 wells of PhenoPlate 384-well microplate (Revvity)). On day 9, the iPSC-microglia were stained with 30 µL of a mixture of NucBlue (1 drop/mL) and CellTracker Deep Red plasma membrane stain (2 µM, Invitrogen, C34565) in Live Cell Imaging Solution (Invitrogen) for 40 minutes at 37°C, 5% CO_2_. Cells were washed once with 50 µL of Hanks Buffered Saline Solution (Invitrogen), and then 50 µL of Microglia media added, with the actin inhibitor cytochalasin D (30 µM for 30 minutes, Tocris, 1233) used in a few wells as a positive control. To initiate phagocytosis, 25 µL of each phagocytic substrate was added to 4 wells per iPSC line, so that a final well concentration of 30 µg/mL E coli bioparticles (Invitrogen), or 133,333 cells/mL pHrodo-SH-SY5Ys, or 1.67 ug/mL pHrodo-myelin was achieved. The plates were incubated for 2 hours at 37°C, 5% CO_2_, and a single z-plane imaged (40x confocal, 1 µm, 10-12 fields per well) in an Opera Phenix at 37°C, 5% CO_2_. Automated image analysis pipelines in Columbus software were used to segment spots of phagocytic substrate within the iPSC-microglia (CellTracker and NucBlue used to identify and segment individual microglia), with medians per well obtained for the total spot area per cell, a parameter chosen because it best reflects both the size and number of solid phagocytosed cargo.

### Statistical analysis

#### Functional phenotypes

All assays were undertaken with replication at multiple levels: 3-10 technical replicates within the same experiment, and 3-4 replicate harvests on consecutive weeks from one differentiation of an iPSC line, and in some cases the same iPSC line was tested in up to 5 independent differentiations. For all assays the median of technical replicates was taken (except for LegendPLEX where mean was used). Harvests were treated as experimental replicates. In any assay that involved counting nuclei, wells with less than 20 nuclei were removed from the analysis. Assay data was then used to construct assay specific models. Importantly, to account for the significant amounts of repeated differentiations associated variation (batch), all models included batch as a random effect in a generalized linear mixed effects model (GLMM). When assays included LPS stimulation, an interaction effect between LPS and polygenic risk (LRWELL/HRLOAD) was included in the model. Each final model was therefore fit using the formula: assay data (continuous) ∼ polygenic risk (binary factor) + stimulation (binary factor) + polygenic risk:stimulation (interaction) + (1|Batch) (random effect). When undertaking comparisons involving *APOE ε4* status, *APOE ε4* status was included as a fixed effect in the model as a factor denoted as either *APOE ε*33 or *APOE ε*34/44, i.e. possessing at least 1 *APOE ε4* allele. The *p*-values for each parameter in the model were then extracted.

When statistically significant interaction effects were found, post-hoc testing to extract individual group comparisons (i.e. between the four combinations of HRLOAD/LRWELL and LPS+/LPS-) were made using estimated marginal means (emmeans function in R). Effect sizes were reported as the percentage difference between the median values of two phenotypes. Given the extensive number of 44 phenotypic parameters (see Supplementary Table 2) undertaken in this work, to reduce the number of false positive findings, we corrected all nominal *p*-values using the Benjamini-Hochberg procedure. A list of all nominal and corrected *p*-values and their assay of origin can be found in Supplementary Table 2. With the exception of plots stratified by *APOE ε4* status, only comparisons that were statistically significant after *p*-value correction were displayed on plots to improve visual clarity. For the ATP fold-change analysis, the median ATP production of individual ATP measurements per iPSC line under stimulated conditions was divided by that under unstimulated conditions. Fold change was then modelled as the outcome variable with genotype as a predictor variable. Fold-change analysis was not performed for cytokine assays given the inflation in fold change caused by near zero values of cytokine produced under unstimulated conditions.

As a secondary exploratory analysis, correlations between assay data that showed a statistically significant or trending effect of genotype was undertaken to understand whether observed effects were independent of one another. First, data points from distinct assays (e.g. mitoATP production and IL-6 production) were matched based on iPSC line and differentiation data, ensuring that matched data points came from the same week since microglial differentiation of the same iPSC line. Pearson correlation was computed for each pair of assay datasets using pairwise complete observations, A Bonferroni correction was applied to this secondary analysis given that a total of 28 pairwise correlations were tested in this secondary analysis. Any correlation with a *p*-value less than 0.05 after correction was denoted as significant, displayed in the correlation matrix and further visualised as grouped scatter plot.

All data were visualised using the *ggplot2* package in R, with further annotations from *ggpubr*, *guppy* and other *tidyverse* extension packages.

#### Proteomics

To account for the repeated measures design, cell line donor was modeled as a random effect by generating a donor-level blocking factor using the duplicateCorrelation() function. Our main statistical model used an interaction design ∼Batch + AD_Risk * Stimulation_Condition to assess the effect of AD polygenic risk at baseline, with LPS stimulation and to ascertain the effect of LPS stimulation in HRLOAD vs LRWELL. Significance was defined using a combination of Benjamini-Hochberg corrected *p*-values (FDR) and a fold-change threshold of 0.2.

To identify proteomic signatures associated with the independent variables, we performed Fast Gene Set Enrichment Analysis (FGSEA) across both the full proteome and functional extremes (amyloid-β, MitoATP, and IL-6). We categorized samples into ‘high’ and ‘low’ groups based on the assay scores. Proteins were ranked for GSEA using a composite score of *p*-value and log fold-change to prioritize proteins with both high effect size and statistical confidence. To ensure biological diversity in the reported results and minimize term redundancy, we implemented a Jaccard similarity filter (maximum 40% gene overlap). This allowed for the selection of the top 10 most robust and biologically distinct signals for each phenotype.

## Data and code availability

Raw mass spectrometry data for the proteomics will be deposited in PRIDE. Code for the statistical analysis and proteomics analysis, and proteomics analysis objects will be available in GitHub from publication: https://github.com/UKDRI/IPMAR_Paper_2026/

## Acknowledgements

The authors would like to thank those involved in the generation and QC of the iPSC lines, previously described in (Maguire et al., 2025), but in particular Sarah Ellwood and Alex Evans. The authors would also like to acknowledge members of the Cardiff DRI analyst group for helpful discussions.

## Funding

This research was supported by funding from the UK Dementia Research Institute (award number UK DRI-3201) through UK DRI Ltd, principally funded by the Medical Research Council, and also The Moondance Foundation.

## Author contributions

Ju.W. directed the overall strategy. Ju W, C.W., P.R.T., V.E-P., N.A., S.A.C., and H.H-R. conceptualised the study. H.H-R., E.M., B.S., R.O., N.C.R., P.R.T., R.S., P.H. and Ju.W. designed and interpreted the experiments. H.H-R., E.M., B.S., R.O., C.B., Ji.W., and N.S. performed cell experiments and analysis of the raw data. R.S. and Ju. W. provided samples from cohort. B.G. and K.F. performed the mass spectrometry and peptide identification. L.C., S.K., R.M, A.C.M., M.B-H., and Ju. W. and P.H. performed statistical analysis of the data and advised on statistical analysis methods. L.C., R.M., C.B., R.O., and H.H-R made the figures. H.H-R., Ju.W., and E.M. wrote the manuscript with input from the other authors.

## Competing interests

S.A.C. has received research funding from GSK, Eli Lilly and Janssen. The remaining authors declare no competing interests.

## Extended data figures

**Extended Data Figure 1.**
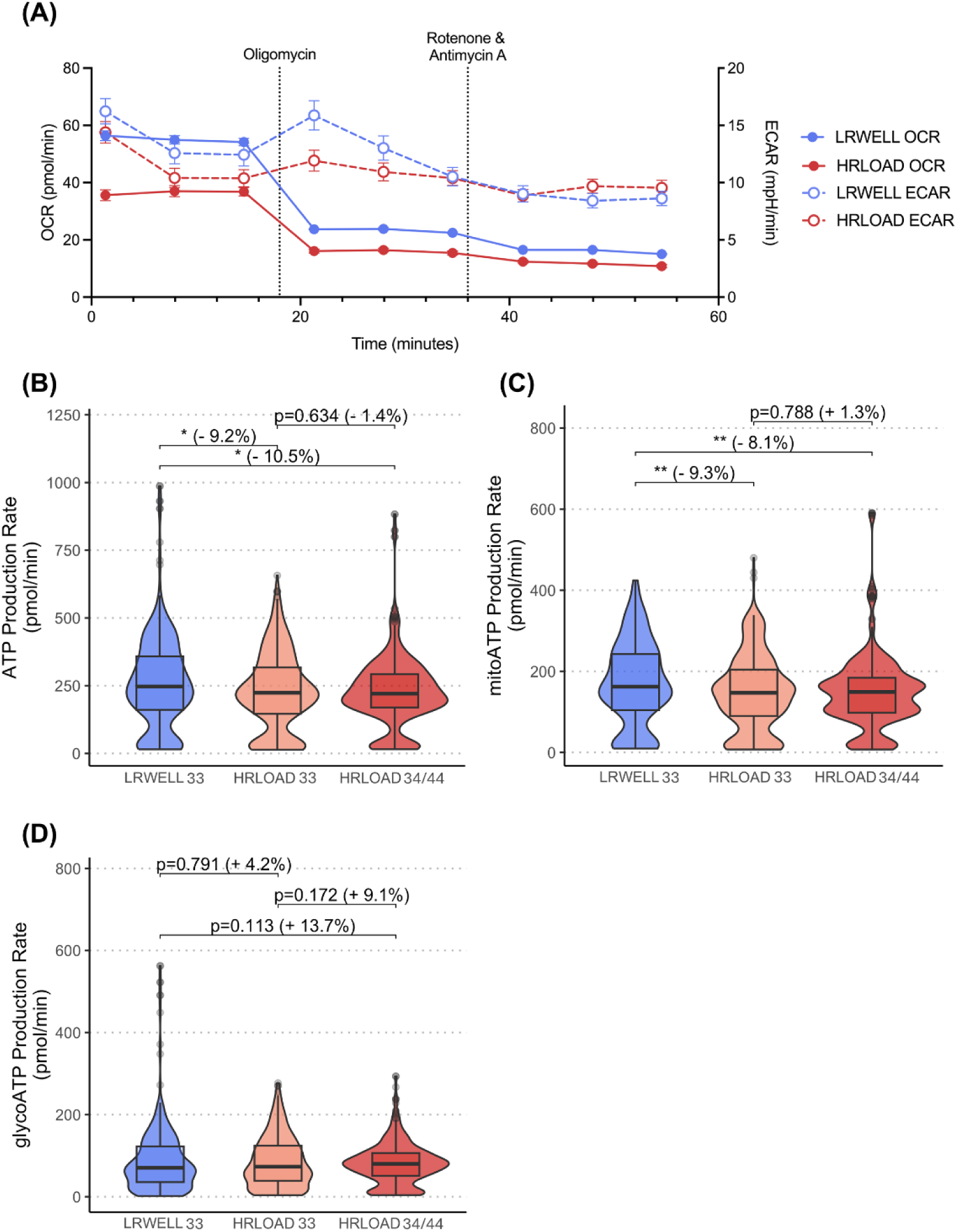
Mitochondrial metabolic dysfunction in HRLOAD microglia stratified by APOE genotype. **(A)** Representative ECAR and OCR trace for LPS stimulated LRWELL (DRICUi020-A) and HRLOAD (DRICUi006-A) iPSC-microglia in the Seahorse XF Real-Time ATP Rate assay. Data points show mean ± SEM of n=5-6 technical replicate wells from 1 assay. **(B)** Total, **(C)** mitochondrial (mitoATP), and **(D)** glycolytic (glycoATP) ATP production rates (pmol/min) were measured using a Seahorse XF Real-Time ATP Rate assay, with LPS stimulation, showing that APOE ε4 does not alter the HRLOAD phenotype. Data plotted is medians obtained from 3-10 replicate wells. Across all parts of the figure, box plots show the median and interquartile range for N= 17 HRLOAD APOE ε33, 17 HRLOAD APOE ε34/44, 17 LRWELL from a minimum of 4 experiments. LRWELL (blue), HRLOAD33 (orange), and HRLOAD34/44 (red) were compared using linear mixed effects modelling, with APOE as a fixed effect. Uncorrected p-values are displayed. * p < 0.05, ** p < 0.01.

**Extended Data Figure 2.**
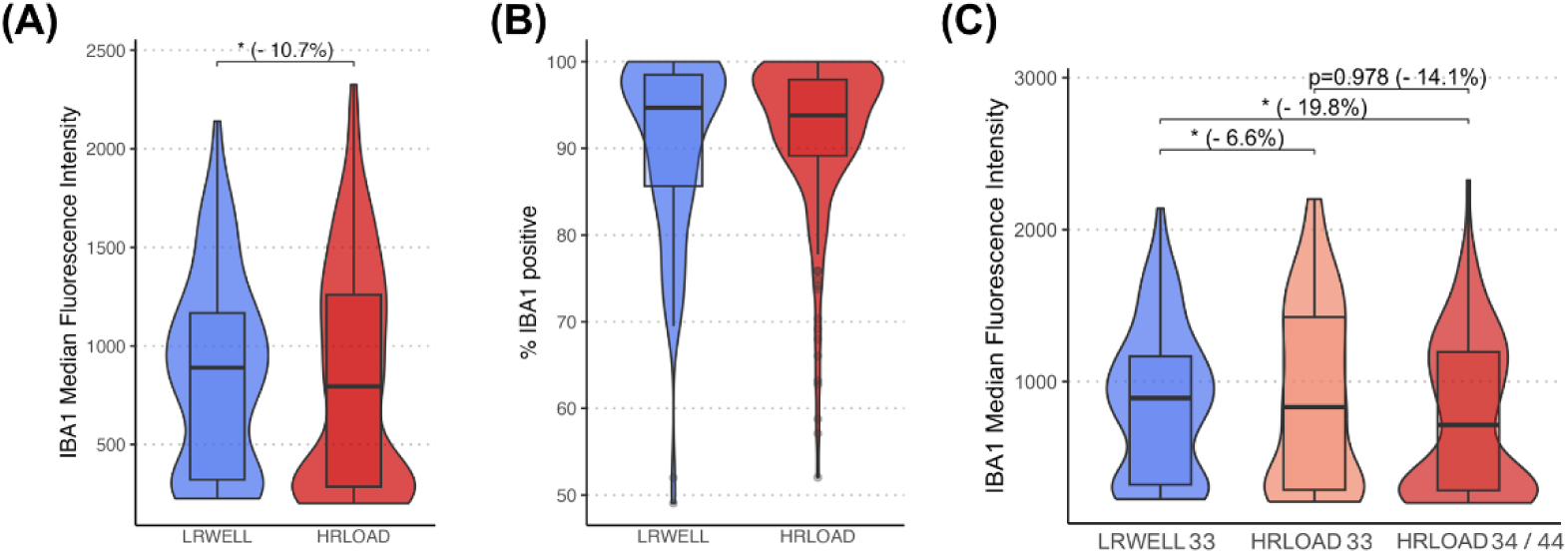
IBA1 expression is reduced in HRLOAD microglia. The microglia activation marker IBA1 was analysed by immunocytochemistry. There was no difference in **(B)** percentage of IBA1+ cells, however a significant difference in **(A)** median fluorescence intensity was found between HRLOAD and LRWELL. Data plotted is medians obtained from 4 replicate wells. For (A and B), box plots show the median and interquartile range for N= 35 HRLOAD, 17 LRWELL from a minimum of 4 experiments. HRLOAD was compared to LRWELL using linear mixed effects modelling with p-values corrected by Benjamini-Hochberg procedure. In HRLOAD iPSC-microglia segregated by APOEε4 status, the IBA1 median fluorescence intensity **(C)** was not reduced in HRLOAD APOE 34/44 versus HRLOAD APOE 33. Data plotted is medians obtained from 4 replicate wells. Box plots show the median and interquartile range for N= 18 HRLOAD APOE ε33, 16 HRLOAD APOE ε34/44, 15 LRWELL from a minimum of 4 experiments. LRWELL (blue), HRLOAD33 (orange), and HRLOAD34/44 (red) were compared using linear mixed effects modelling, with APOE as a fixed effect. Uncorrected p-values are displayed. * p < 0.05.

**Extended Data Figure 3.**
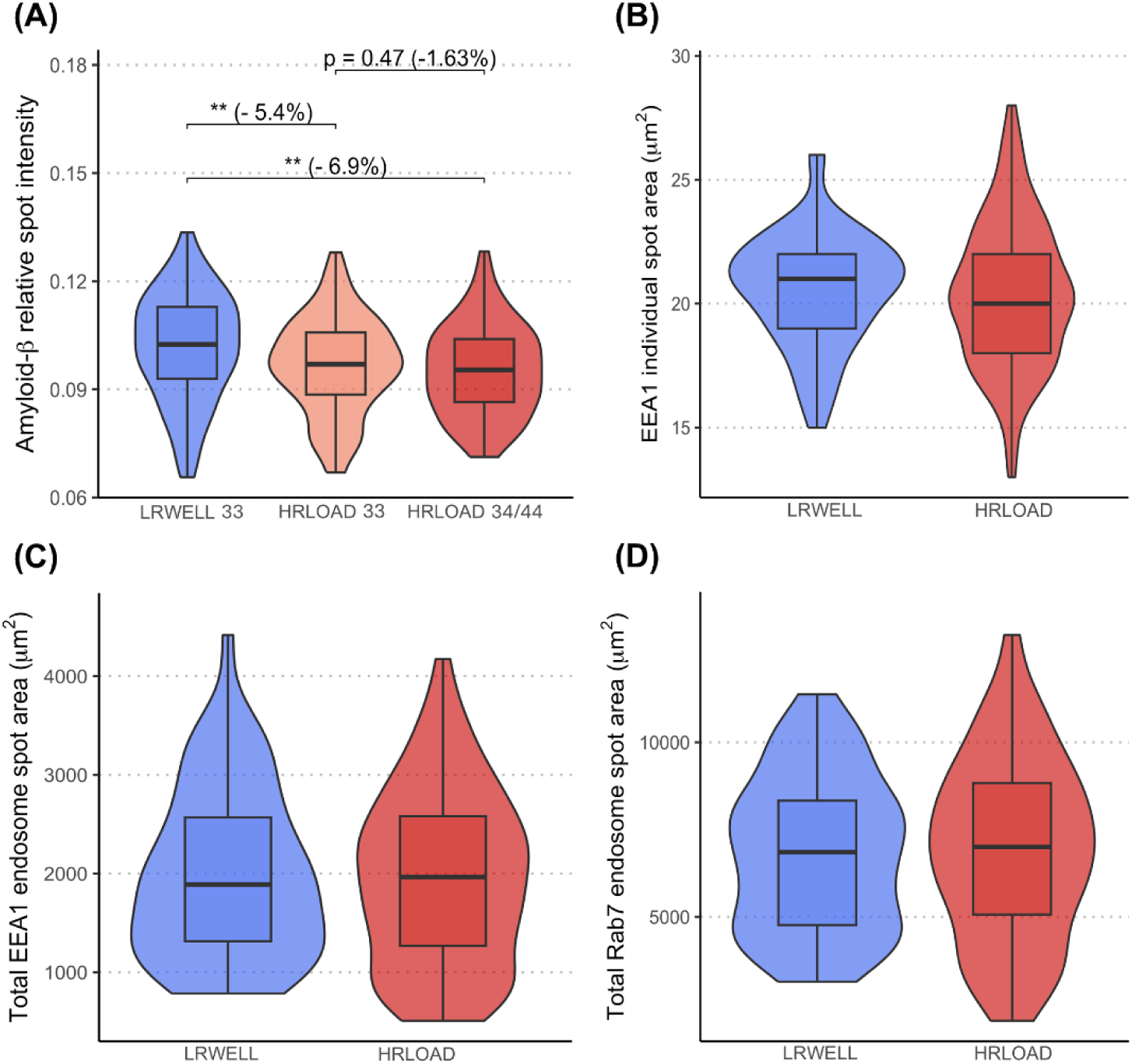
APOE4 does not alter amyloid-β endocytosis. (**A**) In HRLOAD iPSC-microglia segregated by APOEε4 status, i.e. heterozygous for APOE ε3 (HRLOAD APOE 33) or possessing one or two APOEε4 alleles (HRLOAD APOE 34/44), amyloid-β endocytosis were not significantly altered in HRLOAD APOE 34/44 versus HRLOAD APOE 33. Data plotted is medians obtained from 4 replicate wells. Box plots show the median and interquartile range for N= 18 HRLOAD APOE ε33, 16 HRLOAD APOE ε34/44, 15 LRWELL. LRWELL (blue), HRLOAD33 (orange), and HRLOAD34/44 (red) were compared using linear mixed effects modelling, with APOE as a fixed effect. Uncorrected p-values are displayed. Endosome size and quantity are unaltered in HRLOAD microglia. Endosomes were visualized by immunocytochemistry with EEA1 as a marker for early endosomes and Rab7 as a marker for late endosomes. EEA1+ endosome size **(B)** and quantity **(C)** were analysed per cell, and Rab7+ endosome quantity **(D)** per cell. Neither were significantly altered in HRLOAD microglia versus LRWELL. Data plotted is medians obtained from 4 replicate wells. Box plots show the median and interquartile range for N= 23 HRLOAD, 11 LRWELL.. HRLOAD was compared to LRWELL using linear mixed effects modelling with p-values corrected by Benjamini-Hochberg procedure. ** p < 0.01.

**Extended Data Figure 4.**
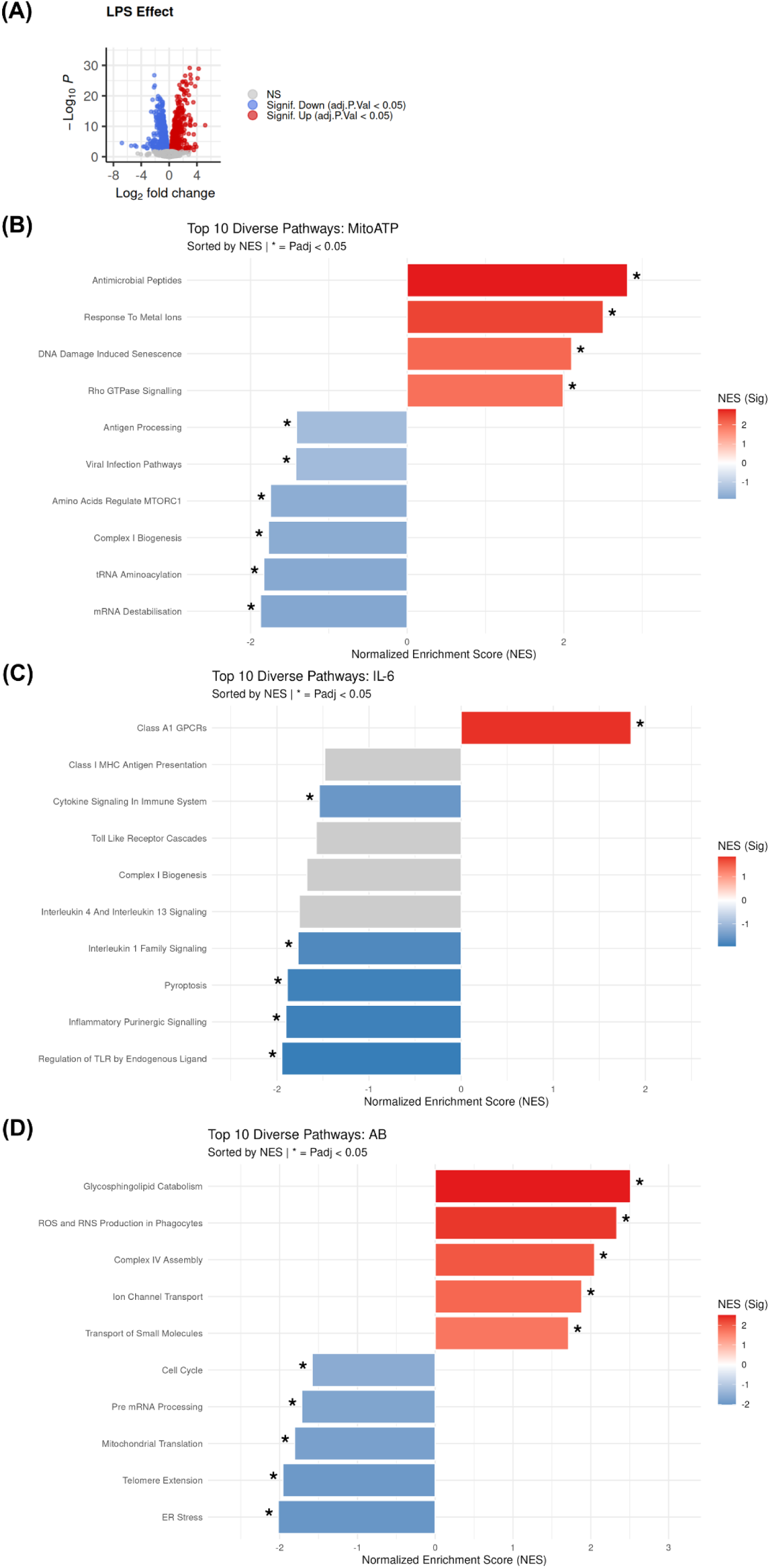
Pathway enrichment of proteomes distinguishing functional extremes. **(A)** Volcano plot of differentially expressed proteins comparing LPS-stimulated to unstimulated, significant upregulated and downregulated proteins are highlighted in red and blue respectively. For the phenotypic assays: **(B)** stimulated Mitochondrial ATP production (MitoATP), **(C)** stimulated IL-6 secretion, **(D)** Amyloid-β (AB) uptake, the top 5 highest and bottom 5 lowest-performing extreme lines were selected and proteins were ranked for GSEA using a composite score of p-value and Log Fold-Change. Bar charts show the most statistically significant Reactome terms filtered by a Jaccard similarity coefficient (cutoff < 0.4) to eliminate redundant gene sets and highlight distinct biological processes. Data is from 2-3 technical replicate wells processed separately from one harvest.

### Supplementary information

- Supplementary Note – containing Supplementary Methods
- Supplementary Table S1 – iPSC used in this study
- Supplementary Table S2 – *p*-values from linear mixed modelling
- Supplementary Table S3 – *p*-values from post-hoc comparisons of significant interactions
- Supplementary Table S4 – Proteomics differential analysis results for unstimulated microglia
- Supplementary Table S5 – Proteomics FGSEA for unstimulated microglia
- Supplementary Table S6 – Proteomics differential analysis results for LPS challenged microglia
- Supplementary Table S7 – Proteomics FGSEA for LPS challenged microglia
- Supplementary Table S8 – Proteomics differential analysis for metabolic extreme (challenged mitochondrial ATP production)
- Supplementary Table S9 – Proteomics FGSEA for metabolic extreme (challenged mitochondrial ATP production)
- Supplementary Table S10 – Proteomics differential analysis for cytokine extreme (challenged IL-6 secretion)
- Supplementary Table S11 – Proteomics FGSEA for cytokine extreme (challenged IL-6 secretion)
- Supplementary Table S12 – Proteomics differential analysis for endocytic extreme (amyloid-β uptake)
- Supplementary Table S13 – Proteomics FGSEA for endocytic extreme (amyloid-β uptake)

## References

Abud, E. M., Ramirez, R. N., Martinez, E. S., Healy, L. M., Cecilia, H. H., Newman, S. A., Yeromin, A. V, Scarfone, V. M., Samuel, E., Fimbres, C., Caraway, C. A., Fote, G. M., Abdullah, M., Agrawal, A., Kayed, R., Gylys, K. H., Cahalan, M. D., Brian, J., Antel, J. P., … Blurton-Jones, M. (2017). iPSC-derived human microglia-like cells to study neurological diseases. Neuron., 94(2), 278–293. 10.1016/j.neuron.2017.03.042

ADGC, Bonn, CHARGE EADB, Bellenguez, C., EADI, FinnGen, GERAD, GR@ACE/DEGESCO, & PGC-ALZ. (2025). Consensus meta-analysis of genome-wide association studies for Alzheimer’s disease and related dementia. MedRxiv (Pre-Print), 2025.10.20.25338060. 10.1101/2025.10.20.25338060

Andreone, B. J., Przybyla, L., Llapashtica, C., Rana, A., Davis, S. S., van Lengerich, B., Lin, K., Shi, J., Mei, Y., Astarita, G., Di Paolo, G., Sandmann, T., Monroe, K. M., & Lewcock, J. W. (2020). Alzheimer’s-associated PLCγ2 is a signaling node required for both TREM2 function and the inflammatory response in human microglia. Nature Neuroscience, 10.1038/s41593-020-0650-6

Bergsbaken, T., Fink, S. L., & Cookson, B. T. (2009). Pyroptosis: host cell death and inflammation. Nature Reviews Microbiology 2009 *7*:2, 7(2), 99–109. 10.1038/nrmicro2070

Cadiz, M. P., Jensen, T. D., Sens, J. P., Zhu, K., Song, W. M., Zhang, B., Ebbert, M., Chang, R., & Fryer, J. D. (2022). Culture shock: microglial heterogeneity, activation, and disrupted single-cell microglial networks in vitro. Molecular Neurodegeneration, 17(1), 26. 10.1186/s13024-022-00531-1

Chen, Z., Balachandran, Y. L., Chong, W. P., & Chan, K. W. Y. (2024). Roles of Cytokines in Alzheimer’s Disease. International Journal of Molecular Sciences, 25(11), 5803. 10.3390/ijms25115803

Codocedo, J. F., Mera-Reina, C., Bor-Chian Lin, P., Fallen, P. B., Puntambekar, S. S., Casali, B. T., Jury-Garfe, N., Martinez, P., Lasagna-Reeves, C. A., & Landreth, G. E. (2024). Therapeutic targeting of immunometabolism reveals a critical reliance on hexokinase 2 dosage for microglial activation and Alzheimer’s progression. Cell Reports, 43(7), 114488. 10.1016/j.celrep.2024.114488

Connolly, N. M. C., Theurey, P., Adam-Vizi, V., Bazan, N. G., Bernardi, P., Bolaños, J. P., Culmsee, C., Dawson, V. L., Deshmukh, M., Duchen, M. R., Düssmann, H., Fiskum, G., Galindo, M. F., Hardingham, G. E., Hardwick, J. M., Jekabsons, M. B., Jonas, E. A., Jordán, J., Lipton, S. A., … Prehn, J. H. M. (2018). Guidelines on experimental methods to assess mitochondrial dysfunction in cellular models of neurodegenerative diseases. Cell Death and Differentiation, 25(3), 542–572. 10.1038/S41418-017-0020-4

De Deyn, L., & Sleegers, K. (2025). The impact of rare genetic variants on Alzheimer disease. Nature Reviews Neurology 2025 21:3, 21(3), 127–139. 10.1038/s41582-025-01062-1

De Strooper, B., Sierksma, A., Moechars, D., Borrie, S., Snellinx, A., Bourdely, P., Thrupp, N., Moonen, S., Pasciuto, E., Humblet-Baron, S., Stevenson-Hoare, J., Simmonds, E., Thal, D., Mazzone, M., Young-Pearse, T., Vandenberghe, R., Escott-Price, V., & Fiers, M. (2025). Polygenic risk for Alzheimer’s disease shapes microglial inflammatory and antigen-presentation programs in vivo. Research Square (Pre-Print). 10.21203/RS.3.RS-7794221/V1

Efthymiou, A. G., & Goate, A. M. (2017). Late onset Alzheimer’s disease genetics implicates microglial pathways in disease risk. Molecular Neurodegeneration, 12(1), 43. 10.1186/s13024-017-0184-x

Escott-Price, V., Sims, R., Bannister, C., Harold, D., Vronskaya, M., Majounie, E., Badarinarayan, N., Morgan, K., Passmore, P., Holmes, C., Powell, J., Brayne, C., Gill, M., Mead, S., Goate, A., Cruchaga, C., Lambert, J. C., Van Duijn, C., Maier, W., … Williams, J. (2015). Common polygenic variation enhances risk prediction for Alzheimer’s disease. Brain, 138(12), 3673. 10.1093/brain/awv268

Fu, J., Wang, R. X., He, J. H., Liu, X. J., Wang, X. X., Yao, J. M., Liu, Y., Ran, C. Z., Ye, Q. S., & He, Y. (2025). Pathogenesis and therapeutic applications of microglia receptors in Alzheimer’s disease. Frontiers in Immunology, 16, 1508023. 10.3389/fimmu.2025.1508023

Gatz, M., Reynolds, C. A., Fratiglioni, L., Johansson, B., Mortimer, J. A., Berg, S., Fiske, A., & Pedersen, N. L. (2006). Role of Genes and Environments for Explaining Alzheimer Disease. Archives of General Psychiatry, 63(2), 168–174. 10.1001/ARCHPSYC.63.2.168

Gawlak-Socka, S., Kowalczyk, E., & Wiktorowska-Owczarek, A. (2026). Unfolded Protein Response at the Crossroads: Integrating Endoplasmic Reticulum Stress with Cellular Stress Networks. International Journal of Molecular Sciences 2026, Vol. 27, 27(4). 10.3390/IJMS27041986

Grootjans, J., Kaser, A., Kaufman, R. J., & Blumberg, R. S. (2016). The unfolded protein response in immunity and inflammation. Nature Reviews Immunology, 16(8), 469–484. 10.1038/NRI.2016.62

Hall-Roberts, H., Agarwal, D., Obst, J., Smith, T. B., Monzón-Sandoval, J., Di Daniel, E., Webber, C., James, W. S., Mead, E., Davis, J. B., & Cowley, S. A. (2020). TREM2 Alzheimer’s variant R47H causes similar transcriptional dysregulation to knockout, yet only subtle functional phenotypes in human iPSC-derived macrophages. Alzheimer’s Research & Therapy, 12, 151. doi10.1186/s13195-020-00709-z

Hall-Roberts, H., Di Daniel, E., James, W. S., Davis, J. B., & Cowley, S. A. (2021). In vitro Quantitative Imaging Assay for Phagocytosis of Dead Neuroblastoma Cells by iPSC-Macrophages. JoVE (Journal of Visualized Experiments), 2021(168), e62217. 10.3791/62217

Huang, Y., Happonen, K. E., Burrola, P. G., O’Connor, C., Hah, N., Huang, L., Nimmerjahn, A., & Lemke, G. (2021). Microglia use TAM receptors to detect and engulf amyloid β plaques. Nature Immunology 2021 22:5, 22(5), 586–594. 10.1038/s41590-021-00913-5

Jenkins, B. C., Neikirk, K., Katti, P., Claypool, S. M., Kirabo, A., McReynolds, M. R., & Hinton, A. (2024). Mitochondria in disease: changes in shapes and dynamics. Trends in Biochemical Sciences, 49(4), 346. 10.1016/j.tibs.2024.01.011

Johnson, E. C. B., Dammer, E. B., Duong, D. M., Ping, L., Zhou, M., Yin, L., Higginbotham, L. A., Guajardo, A., White, B., Troncoso, J. C., Thambisetty, M., Montine, T. J., Lee, E. B., Trojanowski, J. Q., Beach, T. G., Reiman, E. M., Haroutunian, V., Wang, M., Schadt, E., … Seyfried, N. T. (2020). Large-scale Proteomic Analysis of Alzheimer’s Disease Brain and Cerebrospinal Fluid Reveals Early Changes in Energy Metabolism Associated with Microglia and Astrocyte Activation. Nature Medicine, 26(5), 769. 10.1038/s41591-020-0815-6

Jung, E. S., Choi, H., & Mook-Jung, I. (2025). Decoding microglial immunometabolism: a new frontier in Alzheimer’s disease research. Molecular Neurodegeneration 2025 20:1, 20(1), 1–29. 10.1186/S13024-025-00825-0

Kenkhuis, B., Somarakis, A., Kleindouwel, L. R. T., van Roon-Mom, W. M. C., Höllt, T., & van der Weerd, L. (2022). Co-expression patterns of microglia markers Iba1, TMEM119 and P2RY12 in Alzheimer’s disease. Neurobiology of Disease, 167(7829), 105684. 10.1016/j.nbd.2022.105684

Lee, H., Pearse, R. V., Lish, A. M., Pan, C., Augur, Z. M., Terzioglu, G., Gaur, P., Liao, M., Fujita, M., Tio, E. S., Duong, D. M., Felsky, D., Seyfried, N. T., Menon, V., Bennett, D. A., De Jager, P. L., & Young-Pearse, T. L. (2025). Contributions of Genetic Variation in Astrocytes to Cell and Molecular Mechanisms of Risk and Resilience to Late-Onset Alzheimer’s Disease. Glia, 73(6), 1166. 10.1002/glia.24677

Leonenko, G., Baker, E., Stevenson-Hoare, J., Sierksma, A., Fiers, M., Williams, J., de Strooper, B., & Escott-Price, V. (2021). Identifying individuals with high risk of Alzheimer’s disease using polygenic risk scores. Nature Communications, 12(1), 4506. 10.1038/s41467-021-24082-z

Lier, J., Streit, W. J., & Bechmann, I. (2021). Beyond Activation: Characterizing Microglial Functional Phenotypes. Cells, 10(9), 2236. 10.3390/cells10092236

Maguire, E., Connor-Robson, N., Shaw, B., O’Donoghue, R., Stöberl, N., & Hall-Roberts, H. (2022). Assaying Microglia Functions In Vitro. Cells, 11(21), 1–25. 10.3390/cells11213414

Maguire, E., Menzies, G. E., Phillips, T., Sasner, M., Williams, H. M., Czubala, M. A., Evans, N., Cope, E. L., Sims, R., Howell, G. R., Lloyd-Evans, E., Williams, J., Allen, N. D., & Taylor, P. R. (2021). PIP2 depletion and altered endocytosis caused by expression of Alzheimer’s disease-protective variant PLCγ2 R522. The EMBO Journal, 40(17), e105603. 10.15252/embj.2020105603

Maguire, E., Winston, J., Ellwood, S. H., O’Donoghue, R., Shaw, B., Morales, A. C., Keat, S., Evans, A., Marshall, R., Luckcuck, L., Brown, L., Salis, E., Leonenko, G., Denning, N., Allen, N. D., Escott-Price, V., Webber, C., Taylor, P. R., Sims, R., … Hall-Roberts, H. (2025). Modeling common Alzheimer’s disease with high and low polygenic risk in human iPSC: A large-scale research resource. Stem Cell Reports, 20(8), 102570. 10.1016/J.STEMCR.2025.102570

Maninger, J. K., Nowak, K., Goberdhan, S., O’Donoghue, R., & Connor-Robson, N. (2024). Cell type-specific functions of Alzheimer’s disease endocytic risk genes. Philosophical Transactions of the Royal Society B: Biological Sciences, 379(1899). 10.1098/rstb.2022.0378

Mazaheri, F., Snaidero, N., Kleinberger, G., Madore, C., Daria, A., Werner, G., Krasemann, S., Capell, A., Trümbach, D., Wurst, W., Brunner, B., Bultmann, S., Tahirovic, S., Kerschensteiner, M., Misgeld, T., Butovsky, O., & Haass, C. (2017). TREM2 deficiency impairs chemotaxis and microglial responses to neuronal injury. EMBO Reports, 18(7), 1186. 10.15252/EMBR.201743922

Mei, S. Y., Zhang, N., Wang, M. J., Lv, P. R., & Liu, Q. (2024). Microglial purinergic signaling in Alzheimer’s disease. Purinergic Signalling, 21(4), 815. 10.1007/s11302-024-10029-8

Miao, J., Chen, L., Pan, X., Li, L., Zhao, B., & Lan, J. (2023). Microglial Metabolic Reprogramming: Emerging Insights and Therapeutic Strategies in Neurodegenerative Diseases. Cellular and Molecular Neurobiology, 43(7), 3191. 10.1007/S10571-023-01376-Y

Monzón-Sandoval, J., Burlacu, E., Agarwal, D., Handel, A. E., Wei, L., Davis, J., Cowley, S. A., Cader, M. Z., & Webber, C. (2022). Lipopolysaccharide distinctively alters human microglia transcriptomes to resemble microglia from Alzheimer’s disease mouse models. DMM Disease Models and Mechanisms, 15(10). 10.1242/dmm.049349

Muth, C., Hartmann, A., Sepulveda-Falla, D., Glatzel, M., & Krasemann, S. (2019). Phagocytosis of apoptotic cells is specifically upregulated in apoe4 expressing microglia in vitro. Frontiers in Cellular Neuroscience, 13. 10.3389/FNCEL.2019.00181

Pålsson-McDermott, E. M., & O’Neill, L. A. J. (2020). Targeting immunometabolism as an anti-inflammatory strategy. Cell Research, 30(4), 300. 10.1038/s41422-020-0291-z

Perez-Alcantara, M., Washer, S., Chen, Y., Steer, J., Gonzalez-Padilla, D., McWilliam, J., Willé, D., Panousis, N., Kolberg, P., Guerrero, E. N., Alasoo, K., Hall-Roberts, H., Williams, J., Cowley, S. A., Trynka, G., & Bassett, A. (2025). Integrated QTL mapping and CRISPR screening in pooled iPSC-derived microglia reveals genetic drivers of neurodegenerative risk. BioRxiv (Pre-Print), 2025.08.18.670767. 10.1101/2025.08.18.670767

Piers, T. M., Cosker, K., Mallach, A., Johnson, G. T., Guerreiro, R., Hardy, J., & Pocock, J. M. (2019). A locked immunometabolic switch underlies TREM2 R47H loss of function in human iPSC-derived microglia. The FASEB Journal, doi:10.1096/fj.201902447R.

Reitz, C., Rogaeva, E., & Beecham, G. W. (2020). Late-onset vs nonmendelian early-onset Alzheimer disease: A distinction without a difference? Neurology: Genetics, 6(5), e512. 10.1212/NXG.0000000000000512

Samanta, S., Akhter, F., Roy, A., Chen, D., Turner, B., Wang, Y., Clemente, N., Wang, C., Swerdlow, R. H., Battaile, K. P., Lovell, S., Yan, S. F., & Yan, S. S. (2023). New cyclophilin D inhibitor rescues mitochondrial and cognitive function in Alzheimer’s disease. Brain, 147(5), 1710. 10.1093/BRAIN/AWAD432

Sims, R., Hill, M., & Williams, J. (2020). The multiplex model of the genetics of Alzheimer’s disease. Nature Neuroscience, 23(3), 311–322. 10.1038/s41593-020-0599-5

Song, W. M., Joshita, S., Zhou, Y., Ulland, T. K., Gilfillan, S., & Colonna, M. (2018). Humanized TREM2 mice reveal microglia-intrinsic and -extrinsic effects of R47H polymorphism. Journal of Experimental Medicine, 215(3), 745–760. 10.1084/jem.20171529

Tansey, K. E., Cameron, D., & Hill, M. J. (2018). Genetic risk for Alzheimer’s disease is concentrated in specific macrophage and microglial transcriptional networks. Genome Medicine, 10(1), 14. 10.1186/s13073-018-0523-8

Trimmer, P. A., Swerdlow, R. H., Parks, J. K., Keeney, P., Bennett, J. P., Miller, S. W., Davis, R. E., & Parker, W. D. (2000). Abnormal Mitochondrial Morphology in Sporadic Parkinson’s and Alzheimer’s Disease Cybrid Cell Lines. Experimental Neurology, 162(1), 37–50. 10.1006/exnr.2000.7333

Victor, M. B., Leary, N., Luna, X., Meharena, H. S., Scannail, A. N., Bozzelli, P. L., Samaan, G., Murdock, M. H., von Maydell, D., Effenberger, A. H., Cerit, O., Wen, H. L., Liu, L., Welch, G., Bonner, M., & Tsai, L. H. (2022). Lipid accumulation induced by APOE4 impairs microglial surveillance of neuronal-network activity. Cell Stem Cell, 29(8), 1197–1212.e8. 10.1016/j.stem.2022.07.005

Volpato, V., Smith, J., Sandor, C., Ried, J. S., Baud, A., Handel, A., Newey, S. E., Wessely, F., Attar, M., Whiteley, E., Chintawar, S., Verheyen, A., Barta, T., Lako, M., Armstrong, L., Muschet, C., Artati, A., Cusulin, C., Christensen, K., … Lakics, V. (2018). Reproducibility of Molecular Phenotypes after Long-Term Differentiation to Human iPSC-Derived Neurons: A Multi-Site Omics Study. Stem Cell Reports, 11(4), 897. 10.1016/j.stemcr.2018.08.013

Wu, X., Miller, J. A., Lee, B. T. K., Wang, Y., & Ruedl, C. (2025). Reducing microglial lipid load enhances β amyloid phagocytosis in an Alzheimer’s disease mouse model. Science Advances, 11(6), 6038. 10.1126/sciadv.adq6038

Yuan, P., Condello, C., Keene, C. D., Wang, Y., Bird, T. D., Paul, S. M., Luo, W., Colonna, M., Baddeley, D., & Grutzendler, J. (2016). TREM2 haplodeficiency in mice and humans impairs the microglia barrier function leading to decreased amyloid compaction and severe axonal dystrophy. Neuron, 90(4), 724–739. 10.1016/j.neuron.2016.05.003

