## Supplementary Methods for "New Human IPSC Models of Late-onset Alzheimer’s Disease Polygenic Risk Identify Multiple Impairments of Microglial Function"

**Polygenic Risk Score (PRS) Calculations**

Summary statistics from clinically assessed case-control GWAS studied on Alzheimer’s disease (AD) (*N* = 63,926) were used to generate genetic scores (Kunkle et al., 2019). PRS were generated using PRSice-2 using default P-value thresholds. PRSice-2 utilizes the most common approach for PRS calculation of clumping and thresholding (C + T), where markers most strongly associated with the phenotype of interest are preferentially retained. Clumping was performed using an linkage disequilibrium (LD) r^2^=0.1 and a window size of 1000 kb. Incorporating directly genotyped APOE isoforms ε2 and ε4 as separate terms in a regression model with the PRS excluding the APOE region, has been shown to provide higher prediction accuracy than modelling the APOE region as part of the PRS (Leonenko et al., 2021). PRS was calculated excluding the *APOE* region (chromosome 19:44.4–46.5 Mb) due to the high LD in this region (PRS.no.APOE). Subsequently, AD PRS was calculated as a weighted sum of PRS.no.APOE and *APOE*(ε2 + ε4), where *APOE* effects were weighted with effect sizes (B(ε2) = −0.47 and B(ε4) = 1.12). This means that, for APOE ε3 homozygous individuals, their AD PRS is equivalent to their PRS.no.APOE. AD PRS was then adjusted by regressing against 8 principal components to control for population stratification, and then standardised within the sample. Subsequently, AD PRS of all of the individuals in our cohorts were standardised against the mean and standard deviation of the AD PRS of unaffected individuals part of the in 1958 Birth Cohort, as a way to express the PRS as standard deviations against the average risk in population controls.

**Generated of iMG from iPSC- detailed method**

iPSC-microglia (iMG) were differentiated from 6 batches of iPSC lines, with three “batch controls” present in every differentiation, using a previously published protocol with minor modifications (Washer et al., 2022). In brief, embryoid bodies (EB) were set up on 2-3 consecutive days per differentiation, with no more than 4 iPSC lines handled by a person at the same time. Immediately prior to use, an AggreWell-800 plate (STEMCELL Technologies) was firstly prepared: 0.5 mL of Anti-Adherence Rinsing Solution (STEMCELL Technologies) was added to the required wells, centrifuged at 3000×g for 3 minutes, washed briefly with 1 mL PBS before the addition of 1 mL of 2× concentrated EB media (1× concentrated EB media: mTESR-Plus supplemented with 50 ng/mL BMP4 (PeproTech), 50 ng/mL VEGF (PeproTech), and 20 ng/mL SCF (PeproTech)), plus 10 µM Y-27632 (Tocris), and the plate warmed to 37 °C in a cell culture incubator before cell addition. iPSC were lifted at 70-80% confluency by washing with PBS and incubation with 1 mL TrypLE (Gibco) for 5-7 minutes at 37 °C, 5% CO_2_. The cells were diluted 1:8 in PBS, a cell count performed, and the cells pelleted by centrifugation at 400×g for 5 minutes. Cells were resuspended in mTeSR-Plus at a concentration of 3.5×10^6^ cells/mL and 1 mL of iPSC added to the prepared AggreWell-800 plate, which was centrifuged at 100×g for 3 minutes with no braking. The AggreWells were fed daily by a 75% media change (two 50% media changes) with 1× concentrated EB media for 2 days in the morning. On day 2, a few hours after feeding, the EBs were gently transferred into 3 wells of a 6-well low-attachment plate using a 5 mL stripette, ensuring similar numbers of EBs per well, and an additional 3-5 mL of 1× concentrated EB medium added per well. The plates were incubated without disturbance for 4 days, then the media aspirated gently, and the EBs from 3 wells collected together and transferred into two T175 flasks containing 20 mL each of Factory media (X-VIVO 15 (Lonza) supplemented with 1× GlutaMax (Gibco), 50 µM 2-mercaptoethanol (Gibco), 1× penicillin/streptomycin (Gibco), 100 ng/mL M-CSF (PeproTech), 25 ng/mL IL-3 (PeproTech)), and incubated at 37 °C, 5% CO_2_ without disturbance for a week. The flasks, known as “Factories” were fed every 7 days with Factory media after set-up, with care taken to avoid disturbing or aspirating the EBs, and free-floating microglia precursors appeared continuously from week 2 onwards. The following regime of feeding was followed for each differentiation, except extra media added to flasks if microglia precursor production was extremely high. Week 1: +10 mL; weeks 2-4 (harvests 1-3): -/+10-30 mL media (discarded the harvested cells as they are too immature); week 5: +20-30 mL; weeks 6-10 (harvests 4-8): -/+20-50 mL media and the harvested cells were used for experiments. Harvested microglia precursors were passed through a 40µm cell strainer to remove clumps, pelleted by centrifugation at 400×g for 5 minutes, resuspended in Microglia media (Advanced DMEM/F12 (Gibco) supplemented with 1× GlutaMax (Gibco), 1× penicillin/streptomycin (Gibco), 100 ng/mL IL-34 (Peprotech), 10 ng/mL GM-CSF (Gibco), 25 ng/mL M-CSF (Peprotech), 50 ng/mL TGF-β1 (Peprotech)) and counted. Microglia precursors were plated for their final maturation to microglia in plates pre-coated with 10 µg/mL (approximately 1 µg/cm^2^) human plasma fibronectin (Sigma) in PBS for a minimum of 30 minutes, which was aspirated (no washing) immediately before addition of media and cells. Microglia were incubated at 37 °C, 5% CO_2_ for 8-10 days for their final maturation, with a media addition at 7 days for 96-well and 12-well plate formats. Proteomics was performed only on harvest 4. Cell phenotypic assays were performed on four harvests, usually harvests 5-8.

**Mass spectrometry**

The entirety of protein samples were digested using the “SP3” methodology as detailed in (Hughes et al., 2018). Samples were processed using the Bravo automated liquid handling platform (Agilent, USA) adapting the published method files from (Müller et al., 2020). Briefly, samples were alkylated with Tris(2-carboxyethyl)phosphine (TCEP) at 56° for 30 minutes before alkylation for 30 minutes using Iodoacetamide in the dark. Magnetic MagReSyn® Hydroxyl beads (BioResyn) were added to the samples and using acetonitrile and ethanol the proteins were washed prior to digestion. Enzymatic digestion was performed using Trypsin / Lys-C mix (Pierce) using an enzyme to protein ration of 1:20. Digestion was performed at 37 degrees overnight. Digested peptides were eluted from the beads and were lyophilized using a vacuum concentrator (Thermo) before storage at -20° before mass spectrometry analysis.

Peptides were analyzed by injecting an estimated 200 ng onto a Vanquish Neo UHPLC System operating in trap and elute mode coupled with an Orbitrap Astral Mass Spectrometer (Thermo Fisher Scientific). Peptides were loaded onto a PepMap Neo Trap Cartridge (Thermo Fisher Scientific #174500) and analyzed on a C18 EASY-Spray HPLC Column (Thermo Fisher Scientific #ES906) with a 11.8 minute gradient from 1% to 55% Buffer B (Buffer A: 0.1% formic acid in water; Buffer B: 0.08% formic acid in 80:20 acetonitrile:water, 0.7 min at 1.8 µL/min from 1% to 4% B, 0.3 min at 1.8 µL/min from 4% to 8% B, 6.7 min at 1.8 µL/min from 8% to 22.5% B, 3.7 min at 1.8 µL/min from 22.5% to 35% B, 0.4 min at 2.5 µL/min from 35% to 55% B). Eluted peptides were analyzed using data-independent acquisition mode on the mass spectrometer.

Resultant mass spectra files were analysed using DIA-NN v.2.2.1 using the library free method using a human FASTA file (Uniprot, downloaded August 2023). The default parameters were used except for “Heuristic protein inference” and the double pass neural network classifier was used. The DIA-NN Intensity matrix was processed using R (v4.5.1). Raw intensities were log transformed in order to stabilize variance while median normalization was used to account for technical variation between the samples. Technical replicates were combined using the avereps() function in limma (v3.66.0) and proteins detected in less than two samples were removed. Differential protein abundance was modeled using a linear framework.

**References for Supplementary Methods**

Hughes, C. S., Moggridge, S., Müller, T., Sorensen, P. H., Morin, G. B., & Krijgsveld, J. (2018). Single-pot, solid-phase-enhanced sample preparation for proteomics experiments. *Nature Protocols 2018 14:1*, *14*(1), 68–85. https://doi.org/10.1038/s41596-018-0082-x

Kunkle, B. W., Grenier-Boley, B., Sims, R., Bis, J. C., Damotte, V., Naj, A. C., Boland, A., Vronskaya, M., van der Lee, S. J., Amlie-Wolf, A., Bellenguez, C., Frizatti, A., Chouraki, V., Martin, E. R., Sleegers, K., Badarinarayan, N., Jakobsdottir, J., Hamilton-Nelson, K. L., Moreno-Grau, S., … Lieberman, A. P. (2019). Genetic meta-analysis of diagnosed Alzheimer’s disease identifies new risk loci and implicates Aβ, tau, immunity and lipid processing. *Nature Genetics*, *51*(3), 414–430. https://doi.org/10.1038/s41588-019-0358-2

Leonenko, G., Baker, E., Stevenson-Hoare, J., Sierksma, A., Fiers, M., Williams, J., de Strooper, B., & Escott-Price, V. (2021). Identifying individuals with high risk of Alzheimer’s disease using polygenic risk scores. *Nature Communications*, *12*(1), 4506. https://doi.org/10.1038/s41467-021-24082-z

Müller, T., Kalxdorf, M., Longuespée, R., Kazdal, D. N., Stenzinger, A., & Krijgsveld, J. (2020). Automated sample preparation with SP3 for low‐input clinical proteomics. *Molecular Systems Biology 2020 16:1*, *16*(1), MSB199111-. https://doi.org/10.15252/MSB.20199111
